# KCNN4 links PIEZO-dependent mechanotransduction to NLRP3 inflammasome activation

**DOI:** 10.1101/2023.03.08.531717

**Authors:** Li Ran, Tao Ye, Eric Erbs, Stephan Ehl, Nathalie Spassky, Izabela Sumara, Zhirong Zhang, Romeo Ricci

## Abstract

Immune cells sense the microenvironment to fine-tune their inflammatory responses. Patients with cryopyrin associated periodic syndrome (CAPS), caused by mutations in the *NLRP3* gene, develop auto-inflammation triggered by non-antigenic, e.g. environmental cues. However, the underlying mechanisms are poorly understood. Here, we uncover that KCNN4, a calcium-activated potassium channel, links PIEZO-mediated mechanotransduction to NLRP3 inflammasome activation. Yoda1, a PIEZO1 agonist, lowers the threshold for NLRP3 inflammasome activation. PIEZO-mediated sensing of stiffness and shear stress increases NLRP3-dependent inflammation. Myeloid-specific deletion of *PIEZO1/2* protects mice from gouty arthritis. Activation of PIEZO1 triggers calcium influx, which activates KCNN4 to evoke potassium efflux promoting NLRP3 inflammasome activation. Activation of PIEZO signaling is sufficient to activate the inflammasome in cells expressing CAPS-causing NLRP3 mutants via KCNN4. Finally, pharmacologic inhibition of KCNN4 alleviates auto-inflammation in CAPS patient cells and in CAPS-mimicking mice. Thus, PIEZO-dependent mechanical inputs augment inflammation in NLRP3-dependent diseases including CAPS.

**One Sentence Summary:** PIEZO-mediated mechanotransduction stimulates KCNN4-dependent potassium efflux to potentiate NLRP3 inflammasome activation.

## INTRODUCTION

Immune cells primarily respond to pathogen exposure and tissue injury using pattern recognition receptors (PRRs) to elicit an inflammatory response (*1, 2*). However, it is increasingly recognized that other parameters in the microenvironment alter during inflammation and that immune cells are capable to adapt their responses to those variations. More recently, specific changes in immune cell activation and effector function in response to mechanical cues and forces have been put forward (*3*). The process of mechanotransduction governs conversion of mechanical inputs into biochemical or electrical signals activating signaling pathways that ultimately affect cellular function.

Macrophages are particularly sensitive to changes in tissue stiffness or tension as they are adherent and contact-dependent. Canonical macrophage functions such as phagocytosis (*4, 5*), generation of reactive oxygen species (ROS) (*6*), and secretion of cytokines (*4, 7–9*) have been demonstrated to change in response to sensing of substrate rigidity. On the other hand, blood monocytes can respond to high shear stress promoting adhesion, phagocytosis and cytokine secretion (*10*).

Macrophages and monocytes, as other cells, employ conserved mechanisms by which they can sense and respond to multiple types of mechanical forces or to surrounding physical properties of different tissues. Sensing through mechano-gated ion channels recently gained a lot of attention. Among them, PIEZO1 and TRPV4 have been mostly characterized in immune cells. PIEZO1 is a member of the PIEZO family of mechanically activated cation channels (*11*), while TRPV4 represents a member of the TRP superfamily of non-selective cation channels (*12*). In response to mechanical inputs, physical distortion of the plasma membrane directly leads to opening of these channels, triggering ion influx, including Ca^2+^, to elicit signal transduction.

PIEZO1 was shown to co-localize with Toll-like receptor 4 (TLR4) to promote ROS production, thereby enhancing pathogen engulfment and killing upon macrophage activation (*6*). Moreover, a pro-inflammatory macrophage phenotype was shown to be promoted by stiff substrates through PIEZO1-dependent Ca^2+^ influx and NF-κB activation (*7*). PIEZO1 is also activated by cyclical hydrostatic pressure resulting in endothelin 1 (EDN1)-dependent hypoxia-inducible factor 1α (HIF1α) stabilization ensuring efficient clearance of intranasal *Pseudomonas aeruginosa* (*13*). In analogy, the role of TRPV4 has been predominantly investigated in macrophages (*4, 14, 15*).

The inflammasome is a cellular complex that provides a first line of response against pathogens and sterile insults. In most cases, the inflammasome complex consists of a sensing receptor, the adaptor ASC and the effector proteinase Caspase-1. Upon activation, its assembly leads to self-activation of Caspase-1, which cleaves pro-IL-1β and pro-IL-18 into their mature forms promoting their secretion and also cleaves Gasdermin D triggering pro-inflammatory lytic cell death, called pyroptosis (*16–19*). NLRP3 inflammasome activation occurs in two principal steps, (1) priming through Toll-like or cytokine receptor signaling resulting in robust expression of the inflammasome components and (2) activation leading to inflammasome complex assembly, which can be triggered by various stimuli, including a remarkable variety of environmental cues (*16–19*). Gain of function mutations in the *NLRP3* gene lead to the development of Cryopyrin-associated periodic syndromes (CAPS), auto-inflammatory conditions that often occur at joints and eyes, highly mechanical sites, and that can be triggered via cold exposure (*20*). However, whether mechanical inputs are directly linked to NLRP3 inflammasome activation has not been systematically addressed.

In light of a potential role of mechanotransduction in NLRP3-dependent inflammation, we set out to explore whether and how mechanical stress can impact NLRP3 inflammasome activation. We find that activation of PIEZO signaling, either through treatment with agonists, via enhancing of substrate stiffness or via inducing of shear stress, amplifies NLRP3 inflammasome activation. Mechanistically, we provide evidence for PIEZO1 activation to evoke calcium influx to induce KCNN4-dependent potassium efflux, thereby potentiating NLRP3 inflammasome activation. In search for disease relevance of our findings, we also demonstrate this mechanism to contribute to NLRP3-dependent inflammation such as monosodium urate-induced arthritis and CAPS. Hence, our results may pave the way for so far unanticipated therapeutic strategies with the goal to alleviate inflammation in these inflammatory conditions.

## RESULTS

### Activation of PIEZO1 by its agonists potentiates NLRP3 inflammasome activation

To address whether PIEZO-mediated mechanotransduction affects NLRP3 inflammasome activation, we first used Yoda1, a PIEZO1 agonist (*21*), and THP-1 cells, a human acute leukemia monocytic cell line widely used for assessing NLRP3 inflammasome activation. THP-1 cells in suspension were primed with LPS and exposed to nigericin to activate the NLRP3 inflammasome in absence or presence of Yoda1. Yoda1 significantly enhanced susceptibility of THP-1 cells to nigericin-induced inflammasome activation as evidenced by increased amounts of cleaved and secreted IL-1β and Caspase-1 in supernatants measured by ELISA and/or Western blotting (Fig. 1A and 1B) and enhanced pyroptosis determined by Sytox Green uptake (Fig. 1C). Nigericin induces NLRP3 inflammasome activation in a potassium efflux-dependent manner (*22*), we thus wondered if it was the same with potassium efflux-independent activators, imiquimod R837 and its derivative CL097 (*23*). Yoda1 also dramatically lowered the threshold of inflammasome activation in response to R837 or its derivative CL097 (fig. S1A-S1F). No induction of pyroptosis was observed in *NLRP3* knockout (KO) THP-1 cells (Fig. 1C and fig. S1C and S1F) confirming that Yoda1-induced potentiation of inflammasome activation was NLRP3-dependent. The release of cleaved IL-1β and Caspase-1 in response to LPS plus nigericin, R837 or CL097 were also markedly enhanced in Yoda1-stimulated murine bone marrow-derived macrophages (BMDMs) as compared to control-treated cells (fig. S1G-S1L). Nigericin-induced pyroptosis was also significantly amplified by Yoda1 in human peripheral blood mononuclear cells (PBMCs) (Fig. 1D). The PIEZO1 agonists Jedi1 and Jedi2 also enhanced R837-dependent NLRP3 inflammasome activation, although to a lesser extend as compared to Yoda1 (fig. S1M). Both Jedi1 and Jedi2 have been shown to exert weaker and shorter effects on PIEZO1 activation as compared to Yoda1 (*24*). In line with a previous study (*7*), Yoda1 treatment also slightly enhanced the production and release of TNFα (fig. S1G, S1I and S1K). Enhanced TNF production has been primarily linked to PIEZO1-induced NF-kB-dependent transcription of this cytokine (*7*). However, Yoda1 treatment did not affect the expression of NLRP3, pro-Caspase-1, pro-IL1β, ASC, NEK7 and Gasdermin D (Fig. 1B). Formation of ASC specks is a hallmark of cells in which the NLRP3 inflammasome is activated (*25*). LPS-primed BMDMs stimulated with nigericin (Fig. 1E and 1F), R837 (fig. S1N, S1O) or CL097 (fig. S1N and S1O) showed significantly enhanced cell numbers containing ASC specks when cotreated with Yoda1 as compared to cells without Yoda1 treatment. Yoda 1, independent of NLRP3, had no potentiating but, if anything, rather an inhibitory effect on activation of the Pyrin and AIM2 inflammasomes (fig. S2A and S2B).

**Figure 1.**
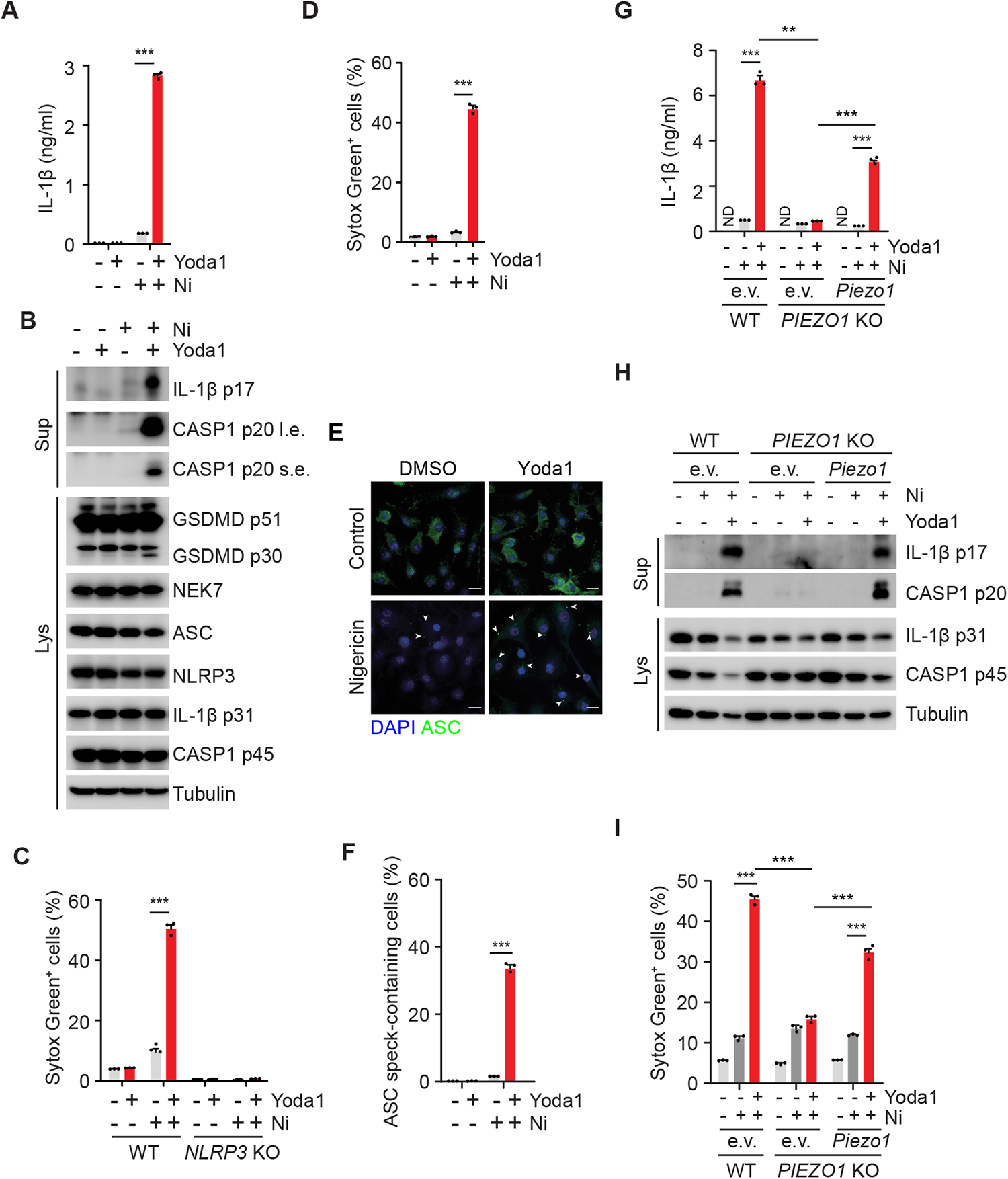
Activation of PIEZO1 by Yoda1 potentiates nigericin-induced NLRP3 inflammasome activation. (**A**) ELISA measurements of IL-1β in culture supernatants from THP-1 cells primed with 1 µg/ml LPS for 3 h and followed by treatment with 5 µM nigericin (Ni) in presence or absence of 25 µM Yoda1. (**B**) Immunoblotting of culture supernatants (Sup) and lysates (Lys) from LPS-primed THP-1 cells treated as in panel A. Antibodies against IL-1β and Caspase-1 (CASP1) were used. An antibody against Tubulin was used as a loading control. “s.e.” short exposure, “l.e.” long exposure. (**C**) Uptake of Sytox Green in LPS-primed wild type (WT) and *NLRP3* KO THP-1 cells treated as described in panel A. (**D**) Uptake of Sytox Green from isolated PBMCs primed with 1 µg/ml LPS for 3 h and followed by treatment with 5 µM nigericin (Ni) in presence or absence of 25 µM Yoda1. (**E**) Representative immunofluorescence images of ASC speck formation in LPS-primed BMDMs stimulated with 2.5 μM nigericin in the presence or absence of 25 µM Yoda1. White arrows indicate ASC specks (green). Scale bars: 10 µm. (**F**) Quantification of macrophages containing ASC specks in experiments as described in panel D. (**G**) ELISA measurements of IL-1β in culture supernatants from WT and *PIEZO1* KO THP-1 cells expressing empty vector (e.v.) or mouse Piezo1. Cells primed with 1 µg/ml LPS for 3 h and followed by treatment with 5 µM nigericin (Ni) in presence or absence of 25 µM Yoda1 for 40 min. (**H**) Immunoblotting of culture supernatants (Sup) and lysates (Lys) from LPS-primed cells treated as described for panel G. Antibodies against IL-1β and Caspase-1 (CASP1) were used. An antibody against Tubulin was used as a loading control. (**I**) Uptake of Sytox Green from LPS-primed cells treated as described in panel F. Sytox Green uptake was analyzed by FACS after staining. “ND” not detected. * *p* < 0.05, ** *p* < 0.01, *** *p* < 0.001. Data shown are representative of at least three independent experiments, except for data shown in panel D which are representative of two independent experiments.

To confirm that the effect of Yoda1 on NLRP3 inflammasome activation was PIEZO-dependent, we next generated *PIEZO1* KO, *PIEZO2* KO and *PIEZO1*/2 double knockout (dKO) THP-1 cells using CRISPR/Cas9-mediated gene editing. Deletion of either gene in THP1 cells was confirmed using Sanger sequencing (fig. S3A-S3C). Yoda1-induced potentiation of inflammasome activation in response to nigericin (fig. S4A-S4C), R837 (fig. S4D-S4F) or CL097 (fig. S4G-S4I) was abolished in *PIEZO1* KO and *PIEZO1/2* dKO cells but not in *PIEZO2* KO cells, confirming that effects of Yoda1 occur through PIEZO1 activation. Neither *PIEZO1* nor *PIEZO2* was required to activate the NLRP3 inflammasome in response to nigericin (fig. S4A-S4C), R837 (fig. S4D-S4F) or CL097 (fig. S4G-S4I) in the absence of Yoda1. Re-expression of PIEZO1 in *PIEZO1* KO THP-1 cells restored increased sensitivity to inflammasome activation in response to nigericin (Fig. 1G-1I), R837 (fig. S5A-S5C) or CL097 (fig. S5D-S5F). To address whether enhancing expression of PIEZO1 potentiates effects of Yoda1 stimulation, we next generated wild-type (WT) and *NLRP3* KO THP-1 cells ectopically expressing PIEZO1. In WT cells ectopically expressing PIEZO1, Yoda1 was sufficient to trigger cleavage and secretion of IL-1β and Caspase-1 as well as pyroptosis in LPS-primed cells, while Yoda1 didn’t show such effects in *NLRP3* KO cells ectopically expressing PIEZO1 (fig. S6A-S6C). Taken together, PIEZO1 activation constitutes a potent mechanism to amplify NLRP3 inflammasome activation in macrophages.

### PIEZO-dependent mechanotransduction modulates NLRP3-dependent inflammation

We next wondered whether physiologic PIEZO activation had an impact on NLRP3 inflammasome activation. To this end, we first subjected THP-1 cells to different cell culture substrate rigidities. THP-1 cells cultured on stiff substrates secreted more IL-1β than cells cultured on soft substrates in response to nigericin and R837 (Fig. 2A). Deletion of both *PIEZO1* and *PIEZO2* attenuated the secretion of IL-1β triggered by nigericin and R837 in cells cultured on stiff as well as on soft substrates (Fig. 2A). We next applied shear stress to cultured THP-1 cells. Likewise, enhanced shear stress amplified IL-1β release in a PIEZO-dependent manner (Fig. 2B). To address whether PIEZO-dependent mechanotransduction was important in NLRP3-dependent inflammation *in vivo*, we next used monosodium urate (MSU) crystal-induced acute arthritis in mice. To this end, intra-articular injections of MSU crystals into ankles of back paws were performed. Ankle swelling over time, and ankle temperature were quantified to evaluate the severity of joint inflammation. To confirm dependence of this model on NLRP3 inflammasome activation, experiments were performed in wild-type (*Nlrp3*^+/+^ mice) and *Nlrp3* knockout mice (*Nlrp3*^-/-^ mice). As expected, ankle swelling, and ankle temperature were significantly attenuated in *Nlrp3*^-/-^ mice as compared to *Nlrp3*^+/+^ mice (Fig. 2C and 2D). We then generated myeloid-specific *Piezo1/2* double knockout mice (*Piezo1^fl/fl^*;*Piezo2^fl/fl^*;*LysM-Cre* mice) by crossing *Piezo1/2* floxed mice (*26, 27*) with lysozyme M (LysM) promoter-dependent Cre recombinase expressing mice (*LysM-Cre* mice) (*28*) and compared them with corresponding *Piezo1/2* floxed control mice (*Piezo1^fl/fl^*;*Piezo2^fl/fl^*). Strikingly, arthritis in mice lacking *Piezo1/2* in the myeloid compartment was significantly attenuated as compared to floxed control mice (Fig. 2E and 2F). These results thus indicate that PIEZO-dependent mechanical inputs are important to modulate NLRP3-dependent inflammation in macrophages *in vitro* and *in vivo*.

**Figure 2.**
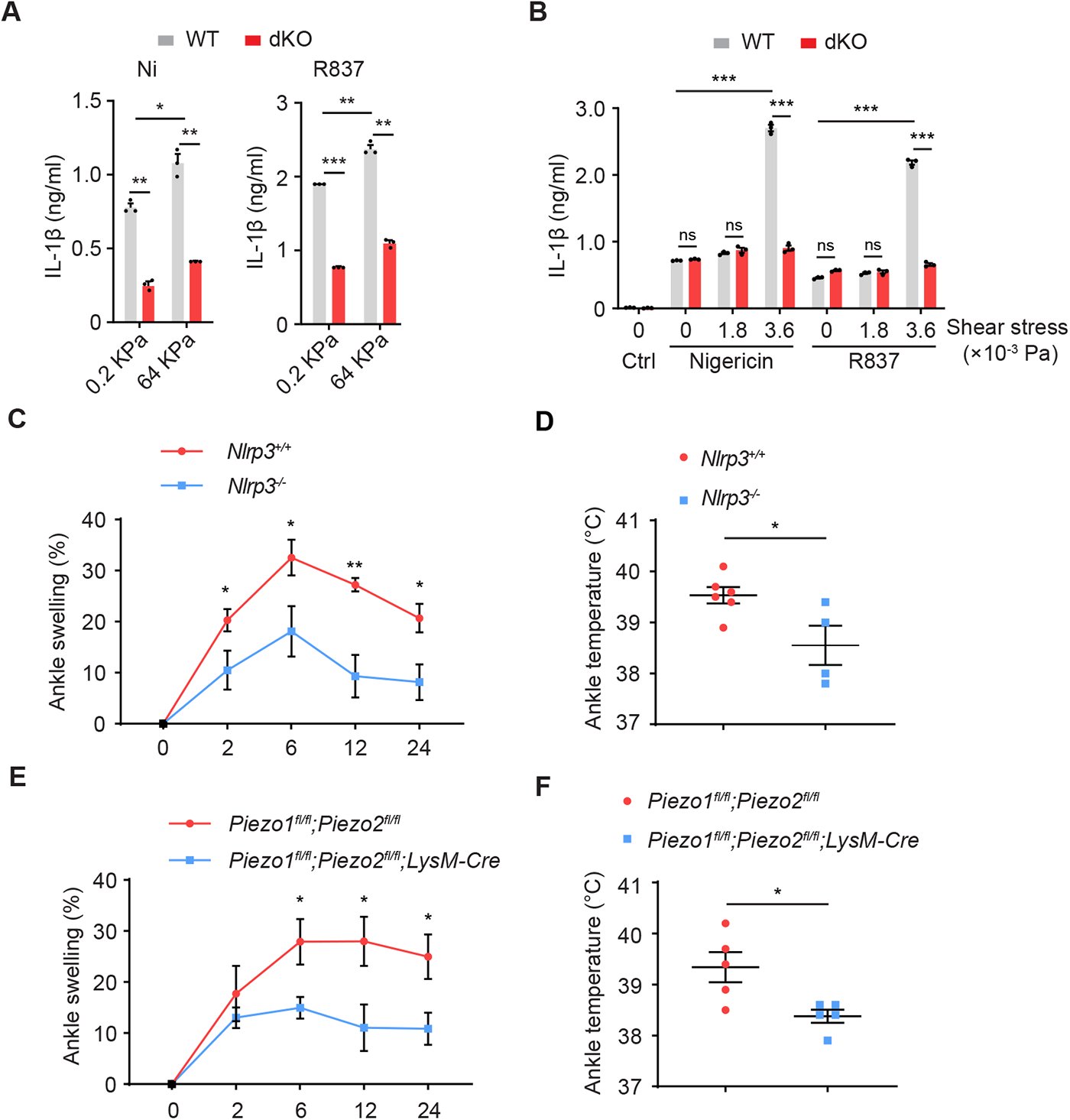
PIEZO-mediated mechanosensing potentiates NLRP3 inflammasome activation. (**A**) ELISA measurements of IL-1β in culture supernatants from WT and *PIEZO1/2* double KO (dKO) THP-1 cells cultured on soft (0.2 KPa) or stiff (64 KPa) silicone. Cells were treated with 5 µM nigericin (Ni) for 40 min or 100 µM R837 for 2 h. (**B**) ELISA measurements of IL-1β in culture supernatants from PMA-differentiated WT and *PIEZO1/2* double KO (dKO) THP-1 cells treated with 5 µM nigericin (Ni) for 40 min or 100 µM R837 for 2 h in absence or presence of shear stress of 1.8×10^-3^ Pa or 3.6×10^-3^ Pa. (**C**) Measurement of the percentage of ankle swelling in *Nlrp3^+/+^* (n=6) and *Nlrp3^-/-^* (n=4) mice at 0, 2, 6 12 and 24 hours after intra-articular injection of MSU (0.5 mg in 10 µl PBS). (**D**) Ankle temperature of *Nlrp3^+/+^*(n=6) and *Nlrp3^-/-^* (n=4) mice at 12 hours after intra-articular injection of MSU. (**E**) Measurement of the percentage of ankle swelling in *Piezo1^fl/fl^;Piezo2^fl/fl^* (n=5) and *Piezo1^fl/fl^;Piezo2^fl/fl^;LysM-Cre* (n=5) mice at 0, 2, 6 12 and 24 hours after intra-articular injection of MSU (0.5 mg in 10 µl PBS). (**F**) Ankle temperature of *Piezo1^fl/fl^;Piezo2^fl/fl^* (n=5) and *Piezo1^fl/fl^;Piezo2^fl/fl^;LysM-Cre* (n=5) mice at 12 hours after intra-articular injection of MSU . “ns” not significant; * *p* < 0.05, ** *p* < 0.01, *** *p* < 0.001. Data shown in panel A are representative of three experiments. Data shown in panel B are representative of two experiments.

### PIEZO-mediated Ca^2+^ influx triggers KCNN4-dependent K^+^ efflux to promote NLRP3 inflammasome activation

Activation of PIEZO channels evoke cellular Ca^2+^ influx (*11*). We thus monitored cytosolic Ca^2+^ level using a genetically-encoded fluorescent Ca^2+^ indicator, jGCaMP7S (*29*), in WT and *PIEZO1/2* dKO THP-1 cells treated with nigericin or R837 in absence or presence of Yoda1. Ionomycin, a potent and selective Ca2+ ionophore, was used as a positive control (Fig. 3A). As previously shown, Yoda1 treatment increased cytosolic Ca^2+^ levels in WT cells. The increase of cytosolic Ca^2+^ levels was further enhanced in response to nigericin (Fig. 3A) or R837 (fig. S7A) in presence of Yoda1. In contrast, cells lacking *PIEZO1* and *PIEZO2* did not show an obvious increase in cytosolic Ca^2+^ in response to Yoda1 or Yoda1 plus nigericin (Fig. 3A) or R837 (fig. S7A). We then asked whether potentiation of NLRP3 inflammasome activation by Yoda1 was Ca^2+^ influx-dependent. To this end, we compared the effect of Yoda1 in THP-1 cells treated in culture medium with or without Ca^2+^. Indeed, enhanced IL-1β and Caspase-1 release as well as pyroptosis in presence of Yoda1 in nigericin-(Fig. 3B-3D), R837-(fig. S7B-S7D) and CL097-(fig. S7E-S7G) treated cells were significantly decreased under Ca^2+^-free medium conditions as compared to cells treated in Ca^2+^-containing medium. Thus, Ca^2+^ influx contributes to PIEZO1-mediated potentiation of NLRP3 inflammasome activation.

**Figure 3.**
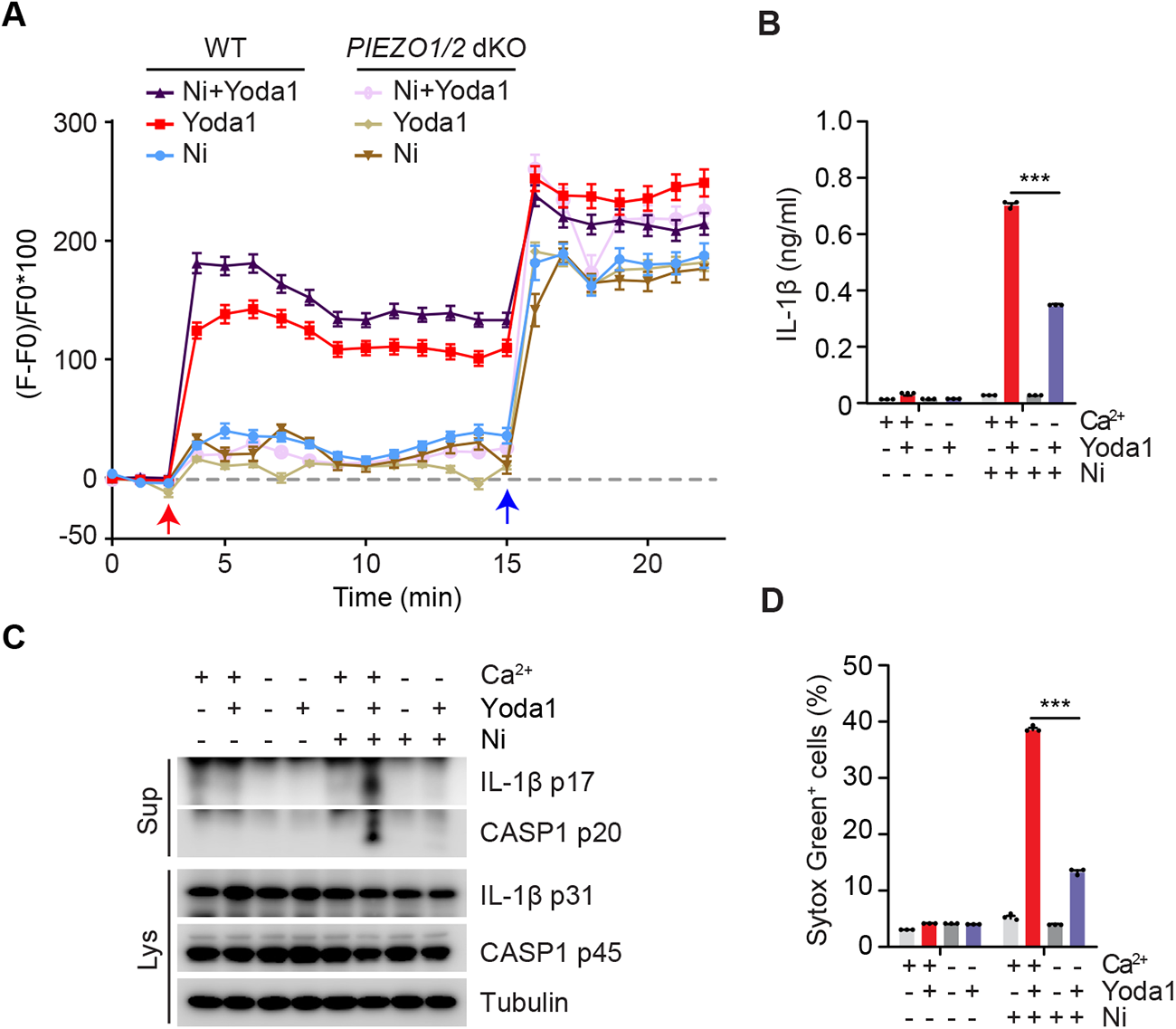
Ca2+ influx is required for PIEZO-mediated mechanosensing and potentiation of NLRP3 inflammasome activation. (**A**) Measurement of cytosolic Ca^2+^ levels in WT and *PIEZO1/2* dKO THP-1 cells using a Ca^2+^ reporter jGCaMP7s. Cells were treated with 5 µM Nigericin (Ni), 25 µM Yoda1 or 5 µM Nigericin plus 25 µM Yoda1 (Ni+Yoda1). The images were acquired using a Nikon spinning-disk microscope with an interval of 60s. Stimuli were added into the culture medium at 3 min as indicated by a red arrow, and ionomycin was added at 15 min as indicated by a blue arrow. Fluorescence intensities of individual cells over time were quantified. n=127 for “WT Ni+Yoda1” group; n=110 for “WT Yoda1” group; n=89 for “WT Ni” group; n=105 for “*PIEZO1/2* dKO Ni+Yoda1” group; n=96 for “*PIEZO1/2* dKO Yoda1” and n=83 for “*PIEZO1/2* dKO Ni” group. (**B**) ELISA measurements of IL-1β in culture supernatants from LPS-primed THP-1 cells treated with 5 µM Nigericin (Ni) in presence or absence of 25 µM Yoda1 in medium with or without (w/o) Ca^2+^. (**C**) Immunoblotting of culture supernatants (Sup) and lysates (Lys) from LPS-primed THP-1 cells in experiments as described for panel B. Antibodies against IL-1β and Caspase-1 (CASP1) were used. An antibody against Tubulin was used as a loading control. (**D**) Uptake of Sytox Green in LPS-primed THP-1 cells in experiments as described for panel B. Sytox Green uptake was analyzed by FACS after staining. ** *p* < 0.01, *** *p* < 0.001. Data are representative of at least three independent experiments.

We next asked how PIEZO1-mediated Ca^2+^ influx amplifies NLRP3 inflammasome activation. To address this question, we performed a genome-wide CRISPR/Cas9-mediated knockout screen in THP-1 cells using the human CRISPR Brunello lentiviral pooled sgRNA library (*30*). R837 was used to induce NLRP3 inflammasome activation to set up this screen. To obtain hits specifically related to PIEZO1-dependent mechanisms of inflammasome activation, we have chosen a R837 concentration in which cotreatment with R837 and Yoda1 led to approximately 85% cell death, while stimulation with R837 alone resulted in basic levels (10 to 15%) of cell death. R837 plus Yoda1-induced cell death was applied twice to ensure maximal elimination of Yoda1-sensitive cells. For the assessment of enrichment of sgRNAs, co-treated cells were compared to vehicle control-treated cells (Fig. 4A). In this screen, NLRP3, ASC (PYCARD) and PIEZO1 were identified as top hits as expected. In addition, ATP11A and CDC50A (TMEM30A), which were reported to directly mediate PIEZO1 activation on membranes (*31*), were also identified. Intriguingly, KCNN4, a Ca^2+^-activated potassium channel, which was reported to act downstream of PIEZO1 in red blood cell (*32*), was the first-ranked hit. We further validated the involvement of KCNN4 in this context using both genetic and pharmacological approaches. Deletion of *KCNN4* using CRISPR/Cas9-mediated gene editing in THP-1 cells (fig. S8A) attenuated the enhancing effect of Yoda1 on R837-induced pyroptosis, cleavage and secretion of IL-1β and Caspase-1 (Fig. 4B-4D and fig. S8B-S8D). Re-expression of EGFP-tagged KCNN4 restored the enhancing effect of Yoda1 on NLRP3 inflammasome activation in *KCNN4* KO cells (Fig. 4B-4D). Deletion of *KCNN4* didn’t affect NLRP3 inflammasome activation induced by R837 alone (Fig. 4B-4D) indicating that KCNN4, as PIEZO proteins, is not required for NLRP3 inflammasome activation in the absence of Yoda1. To further consolidate the role of potassium efflux mediating the effect of Yoda1 on NLRP3 inflammasome activation, we blocked potassium efflux by increasing extracellular potassium concentrations and assessed their effect on NLRP3 inflammasome activation. In line with previous studies (*22, 23, 33*), increasing extracellular potassium concentrations completely blocked nigericin-induced (Fig. 4E-4G), but not R837-induced (fig. S9A and S9B) NLRP3 inflammasome activation. However, increasing extracellular potassium concentrations completely blocked enhancing effects of Yoda1 on R837-induced NLRP3 inflammasome activation (Fig. 4E-4G and fig. S9C-S9G). Consistently, pharmacological inhibition of KCNN4 by TRAM-34 (*34*) similarly inhibited pyroptosis, cleavage and secretion of IL-1β and Caspase-1 induced by R837 plus Yoda1 in THP-1 cells (fig. S9C-S9E). Similar results were also obtained in BMDMs (fig. S9F-S9G). These results suggest that potassium efflux via KCNN4 mediates the enhancing effect of PIEZO1 activation on NLRP3 inflammasome activation.

**Figure 4.**
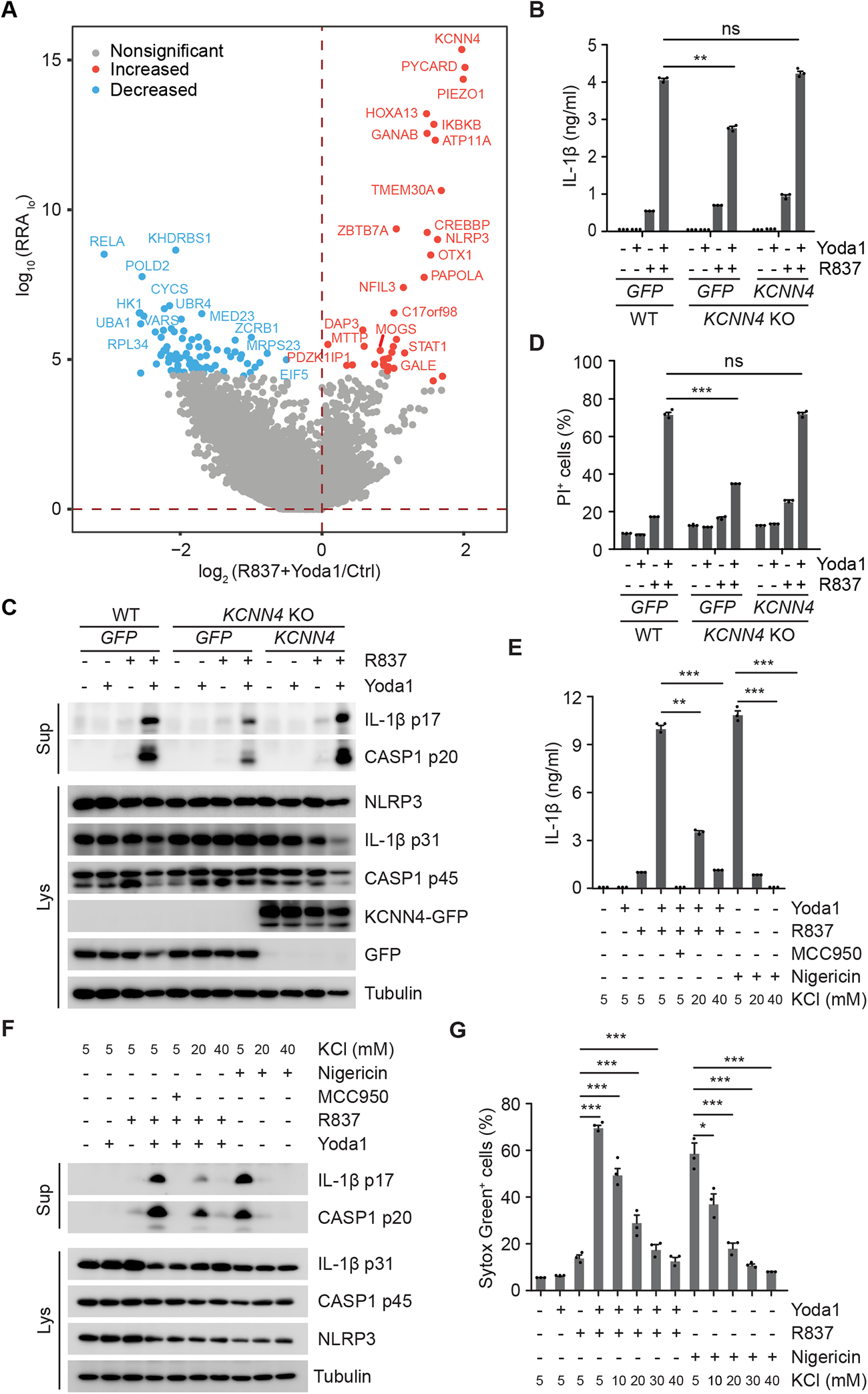
KCNN4 acts downstream of PIEZO-mediated calcium influx to promote NLRP3 inflammasome activation. (**A**) Genome-wide CRISPR/Cas9 screening identifies genes involved in Yoda1-mediated pyroptosis. RRA scores for the comparison of R837 plus Yoda1 vs vehicle. (**B**) ELISA measurements of IL-1β in culture supernatants from WT and *KCNN4* KO THP-1 cells expressing GFP or GFP-tagged KCNN4 (KCNN4). Cells primed with 1 µg/ml LPS for 3 h and followed by treatment with 100 µM R837 in presence or absence of 25 µM Yoda1 for 1 h. (**C**) Immunoblotting of culture supernatants (Sup) and lysates (Lys) from cells in experiments as described in panel B. Antibodies against IL-1β and Caspase-1 (CASP1) were used. An antibody against Tubulin was used as a loading control. (**D**) Uptake of Propidium Iodide (PI) in LPS-primed cells in experiments as described for panel B. (**E**) ELISA measurements of IL-1β in culture supernatants from LPS-primed BMDMs treated with 50 µM R837 or 15 µM nigericin with or without 25 µM Yoda1 in medium containing indicated concentrations of extracellular KCl. (**F**) Immunoblotting of culture supernatants (Sup) and lysates (Lys) from LPS-primed BMDMs in experiments as described for panel E. (**G**) Uptake of Sytox Green in LPS-primed THP-1 cells followed by treatment with 100 µM R837 or 15 µM nigericin with or without 25 µM Yoda1 in medium containing indicated concentrations of extracellular KCl. “ns” not significant, * *p* < 0.05, ** *p* < 0.01, *** *p* < 0.001. Data are representative of at least three independent experiments.

### PIEZO-KCNN4-dependent K^+^ efflux promotes NLRP3-dependent auto-inflammation in CAPS cells

In CAPS patients, auto-activation of NLRP3 inflammasome occurs in absence of infection or tissue injury (*20*). However, inflammation frequently occurs at sites of mechanical impact and in response to cold exposure (*20*). We thus asked whether PIEZO-dependent mechanotransduction was important in triggering auto-activation of the NLRP3 inflammasome in CAPS. To this end, we expressed three CAPS-causing NLRP3 (mouse) mutants, D301N, T346M and R258W, in *NLRP3*-deficient THP-1 cells. *NLRP3*-deficient cells expressing WT NLRP3 or GFP and WT cells expressing GFP were used as controls. Expression of NLRP3 and GFP was confirmed by Western blotting (fig. S10A). Pyroptotic cell death was determined by propidium iodide incorporation. In line with previous studies (*35–38*), cells expressing CAPS-causing NLRP3 mutants displayed significantly enhanced pyroptosis in response to LPS priming alone as compared to control-treated cells (fig. S10B). Importantly, Yoda1 stimulation, similar to LPS treatment, was sufficient to induce pyroptosis in CAPS-causing NLRP3 mutant expressing cells (fig. S10B). Like for LPS treatment, Yoda1 had only minor effects on induction of pyroptosis in control cells (fig. S10B). To verify that Yoda1-induced pyroptosis in CAPS mutant expressing cells was PIEZO-dependent, we next expressed GFP, WT NLRP3 and CAPS-causing NLRP3 mutants in WT, *PIEZO1* KO, *PIEZO2* KO and *PIEZO1/2* dKO THP-1 cells. The expression of NLRP3 and GFP was confirmed by Western blotting (fig. S10C). LPS-induced pyroptosis in cells expressing CAPS-causing NLRP3 mutants was not dependent on PIEZO1 or PIEZO2 as the response was unaltered in *PIEZO1* KO, *PIEZO2* KO and *PIEZO1/2* dKO cells (fig. S10D). However, Yoda1-induced pyroptosis in cells expressing CAPS-causing NLRP3 mutants depended on PIEZO1 expression (Fig. 5A). We further tested whether PIEZO1-dependent activity was KCNN4-dependent. Indeed, inhibition of KCNN4 dramatically inhibited Yoda1-induced inflammasome activation in these cells (Fig. 5B and 5C). To test whether mechanostimulation alone, in the absence of Yoda1, could activate NLRP3 in cells expressing CAPS-causing NLRP3 mutants, we next subjected WT and *PIEZO1/2* dKO THP-1 cells expressing GFP, WT NLRP3 and CAPS-causing NLRP3 mutants to changes in substrate stiffness. Increasing substrate stiffness significantly enhanced IL-1β release in WT cells expressing GFP, WT NLRP3 and mutant NLRP3 expressing THP-1 cells, while this response was significantly reduced in corresponding PIEZO1/2 dKO cells (Fig. 5D). The increase was more pronounced in mutant NLRP3 expressing cells indicating that CAPS mutations render cells more susceptible to changes in substrate stiffness. To corroborate a role of PIEZO-KCNN4-dependent NLRP3 inflammasome activation in cells endogenously expressing a CAPS-causing NLRP3 mutant, we next used mice carrying an intronic floxed neomycin-resistance gene (NeoR that prevents the expression of *Nlrp3 A350V*, a mutation genetically similar to the one observed in human Muckle-Wells syndrome, a distinct subgroup of CAPS (*36*). These mice were crossed with *LysM* promoter-dependent Cre recombinase expressing mice to obtain myeloid-specific expression of mutant NLRP3 (*Nlrp3^A350V/+^; LysM-Cre^+^* mice) as compared to mice lacking Cre recombinase and mutant NLRP3 expression (*Nlrp3^A350V/+^; LysM-Cre^-^* mice). Liver macrophages were isolated from these mice and primed with low dose of LPS to ensure the expression of NLRP3 and IL-1β and further treated with Yoda1. Low dose of LPS induced moderate secretion of IL-1β as expected (Fig. 6A). Treatment of Yoda1 potentiated secretion of IL-1β and Caspase-1, while no IL-1β was detected in cells isolated from control mice under tested conditions (Fig. 6A and 6B). Importantly, inhibition of KCNN4 attenuated Yoda1-induced inflammasome activation in these cells (Fig. 6A and 6B).

**Figure 5.**
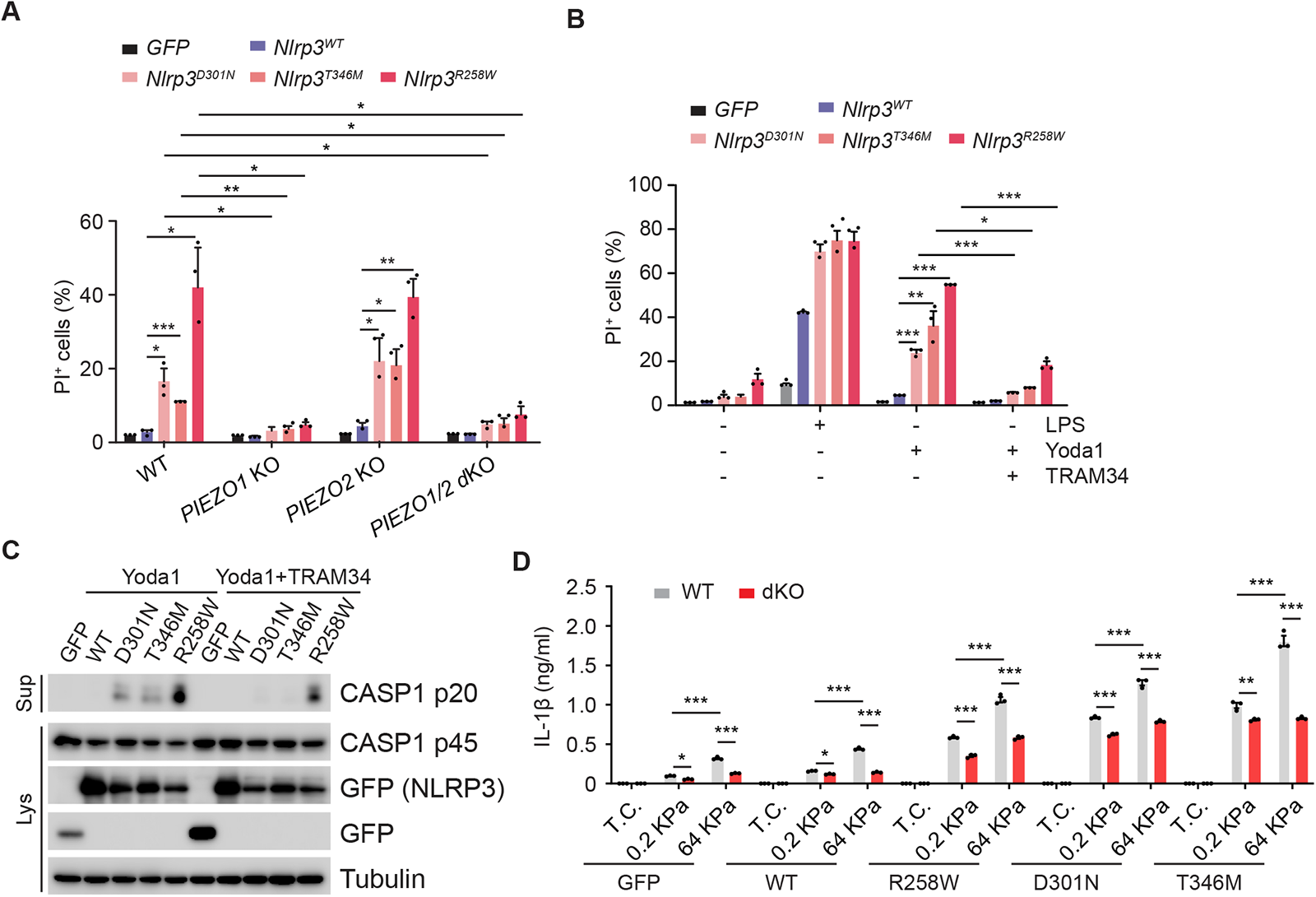
PIEZO stimulation is sufficient to trigger NLRP3 inflammasome activation in cultured cells expressing CAPS-causing NLRP3 mutants. (**A**) Uptake of Propidium Iodide (PI) in WT, *PIEZO1* KO, *PIEZO2* KO and *PIEZO1/2* dKO THP-1 cells expressing GFP, WT or D301N-, T346M-, R258W-mutated mouse Nlrp3. Cells were treated with 25 µM Yoda1 for 6 h. (**B**) Uptake of Propidium Iodide (PI) in WT THP-1 cells expressing GFP, WT or D301N-, T346M-, R258W-mutated mouse Nlrp3. Cells were treated with vehicle, 1 µg/ml LPS, 25 µM Yoda1 or 25 µM Yoda1 plus 5 µM TRAM34 for 6h. (**C**) Immunoblotting of culture supernatants (Sup) and lysates (Lys) from WT THP-1 cells expressing GFP, WT or D301N-, T346M-, R258W-mutated mouse Nlrp3. Cells were treated with 25 µM Yoda1 in presence or absence of 5 µM TRAM34 for 6h. Antibodies against Caspase-1 (CASP1) and GFP were used. An antibody against Tubulin was used as a loading control. (**D**) ELISA measurements of IL-1β in culture supernatants from WT and *PIEZO1/2* dKO (dKO) THP-1 cells expressing GFP, WT or R258W-, D301N-, T346M-mutated mouse Nlrp3 cultured on normal tissue culture plates (T.C.), soft (0.2 KPa) or stiff (64 KPa) silicone. Cells were treated with 100 nM PMA for 3 hours and supernatants were collected at 4 hours after replacement of fresh medium. * *p* < 0.05, ** *p* < 0.01, *** *p* < 0.001. Data are representative of at least three independent experiments.

**Figure 6.**
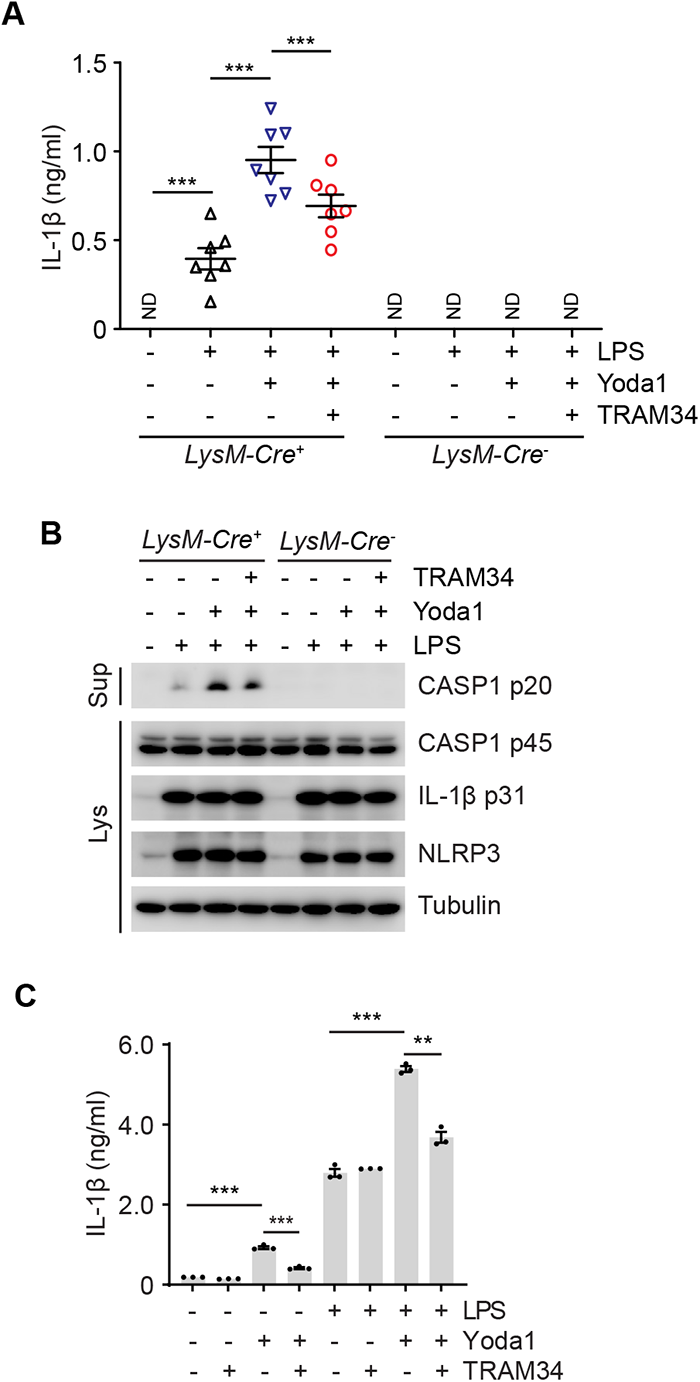
PIEZO1 stimulation is sufficient to trigger NLRP3 inflammasome activation in primary cells isolated from CAPS-mimicking mice and a CAPS patient. (**A**) ELISA measurements of IL-1β in culture supernatants in fetal liver macrophages from *Nlrp3^A350V/+;^LysM-Cre^+^* (*LysM*-Cre^+^) and *Nlrp3^A350V/+^;LysM-Cre^-^*(*LysM-Cre*^-^) mice (n=7 for each genotype) as indicated. Cells were primed with 1 ng/ml LPS for 1 h, followed by treatment with vehicle or 25 µM Yoda1 in presence or absence of 5 µM TRAM34 for 3 h. (**B**) Immunoblotting of culture supernatants (Sup) and lysates (Lys) from cells in experiments as described in panel B. Antibodies against NLRP3, IL-1β and Caspase-1 (CASP1) were used. An antibody against Tubulin was used as a loading control. (**C**) ELISA measurements of IL-1β in culture supernatants from isolated PBMCs of a CAPS patient (*NLRP3 E313K* mutation). Cells were treated with or without 1 µg/ml LPS in presence of vehicle, 25 µM Yoda1, 5 µM TRAM34 or 25 µM Yoda1 plus 5 µM TRAM34 for 4 h. “ND” not detected, * *p* < 0.05, ** *p* < 0.01, *** *p* < 0.001. Data are representative of at least three independent experiments.

To corroborate these findings in primary human cells, we next isolated PBMCs of a CAPS patient carrying a heterozygous germline *NLRP3 E313K* mutation. Even though less pronounced as compared to LPS stimulation, Yoda1 treatment alone was sufficient to enhance secretion of IL-1β from patient PBMCs, which was significantly inhibited upon KCNN4 inhibition (Fig. 6C). Yoda1 also amplified IL-1β in cells co-treated with LPS as compared to cells treated with LPS alone. Inhibition of KCNN4 also inhibited this effect of Yoda1 but had no effect on IL-1β release induced by LPS alone.

Thus, activation of PIEZO1 is sufficient to activate the inflammasome in a KCNN4-dependent manner in cells expressing CAPS-causing NLRP3 mutants.

### Pharmacologic inhibition of KCNN4 activity attenuates CAPS in mice

To open a therapeutic perspective from above findings, we finally asked the question whether pharmacologic KCNN4 inhibition had a beneficial effect on auto-inflammation in CAPS. It was recently reported that macrophage-specific expression of Nlrp3^A350V^ suffices to drive CAPS in mice (*39*). The chemokine receptor Cx3xr1 is widely expressed in the mononuclear phagocyte system. Cre-driven recombination in *Cx3cr1-CreER* mice after tamoxifen induction was reported to occur in tissue-resident macrophages (*40*), likely to be exposed to mechanical stress. Therefore, we crossed *Nlrp3^A350V/A350V^*mutant mice with *Cx3cr1-CreER* mice to obtain *NLRP3^A350V/+^;Cx3cr1-CreER^+^* mice with tamoxifen-inducible expression of NLRP3^A350V^. After tamoxifen induction, *NLRP3^A350V/+^;Cx3cr1-CreER^-^* control mice and *NLRP3^A350V/+^;Cx3cr1-CreER^+^* mice were treated with vehicle or the KCNN4 inhibitor TRAM34 (Fig. 7A). In line with previous studies showing that tamoxifen-induced global Nlrp3^A350V^ expression triggered mild nonlethal autoinflammation associated with body weight loss (*41, 42*), vehicle-treated *NLRP3^A350V/+^;Cx3cr1-CreER*^+^ mice showed gradual body weight loss during treatment; while vehicle-treated *NLRP3^A350V/+^;Cx3cr1-CreER*^-^ control mice didn’t lose their weight (Fig. 7B). Importantly however, TRAM34-treated *Nlrp3^A350V/+^;Cx3cr1-CreER^+^*mice did not experience a significant body weight loss (Fig. 7B). Body weight loss inversely correlated with changes in body temperature (Fig. 7C) and serum levels of IL-1β (Fig. 7D) in mice with corresponding genotypes, indicating that autoinflammation in adult CAPS-mimicking mice is alleviated in response to inhibition of KCNN4.

**Figure 7.**
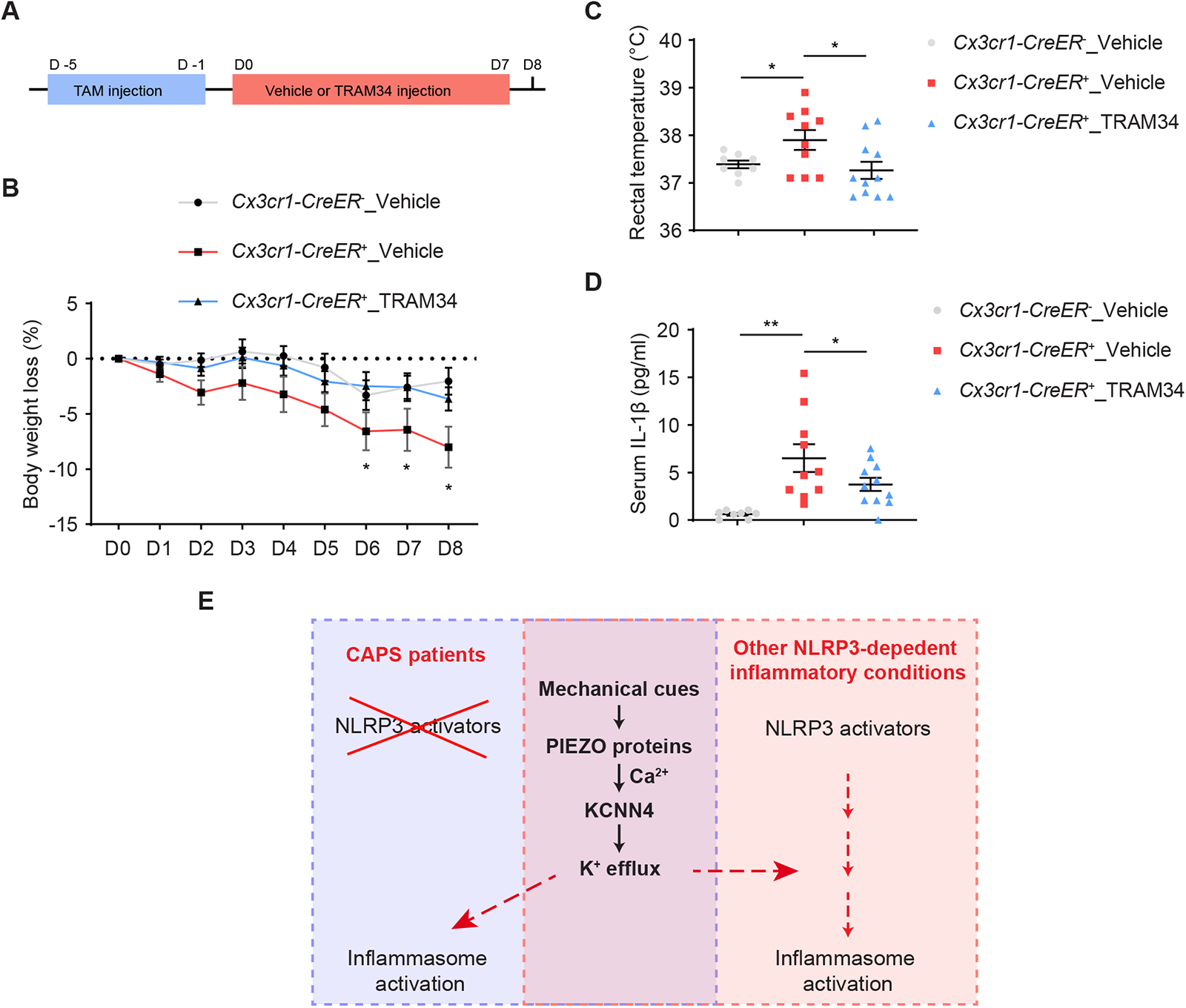
Inhibition of KCNN4 attenuates autoinflammation in CAPS-mimicking mice. (**A**) Experimental timeline. Adult mice were injected intra-peritoneally with 50 mg/kg Tamoxifen (TAM) for five consecutive days and followed by daily intra-peritoneal injections of vehicle or 15 mg/kg TRAM34 for eight consecutive days. On day 8 (D8), rectal temperature was measured and blood was collected. (**B**) Percentage of body weight loss of *Nlrp3^A350V/+;^Cx3cr1-CreER^-^* (*Cx3cr1-CreER*^-^) and *Nlrp3^A350V/+;^Cx3cr1-CreER^+^* (*Cx3cr1-CreER^+^*) mice injected intra-peritoneally with vehicle or 15 mg/kg TRAM34 after tamoxifen induction. (**C**) Rectal temperature at D8 of *Nlrp3^A350V/+;^Cx3cr1-CreER^-^* (*Cx3cr1-CreER*^-^) and *Nlrp3^A350V/+;^Cx3cr1-CreER^+^* (*Cx3cr1-CreER^+^*) mice intra-peritoneally injected with vehicle or 15 mg/kg TRAM34 after tamoxifen induction. (**D**) Serum IL-1β at D8 of *Nlrp3^A350V/+;^Cx3cr1-CreER^-^*(*Cx3cr1-CreER*^-^) and *Nlrp3^A350V/+;^Cx3cr1-CreER^+^* (*Cx3cr1-CreER^+^*) mice intra-peritoneally injected with vehicle or 15 mg/kg TRAM34 after tamoxifen induction. n=8 for *Nlrp3^A350V/+;^Cx3cr1-CreER^-^*treated with vehicle (*Cx3cr1-CreER^-^*_Vehicle); n=10 for *Nlrp3^A350V/+;^Cx3cr1-CreER^+^*mice treated with vehicle (*Cx3cr1-CreER^+^*_Vehicle); n=11 for *Nlrp3^A350V/+;^Cx3cr1-CreER^+^* mice treated with TRAM34 (Cx3cr1-*Cx3cr1-CreER^+^*_TRAM34). * *p* < 0.05, ** *p* < 0.01. (**E**) Proposed model of the mechanism linking mechanostimulation to NLRP3 inflammasome activation *in vivo*.

Altogether, our results suggest that KCNN4 acts downstream of PIEZO signaling-induced Ca^2+^ influx to promote NLRP3 inflammasome activation, a mechanism that may contribute to microenvironmental effects on severity of CAPS and other NLRP3-dependent inflammatory conditions.

## DISCUSSION

Immune cells including macrophages are constantly exposed to microenvironmental changes, such as shear stress in the circulation during extravasation, membrane reorganization during transmigration, hydrostatic pressure in blood vessels, lung and heart (*27*). These changes redirect their responses to physiological and pathological cues (*27*). The NLRP3 inflammasome plays critical roles in innate immune responses detecting various endogenous and exogenous danger signals. In this study, we have provided a direct link between PIEZO protein-mediated mechanosensing and the NLRP3 inflammasome-dependent immune response. Based on our data, we suggest a model, whereby mechanical cues act in parallel to trigger KCNN4-dependent K^+^ efflux (Fig. 7E). In CAPS, a condition of sterile auto-inflammation, no specific NLRP3 activator is present. Therefore, KCNN4-dependent K^+^ efflux might represent a key step to induce NLRP3 inflammasome activation. This is supported by our data showing that PIEZO activation is sufficient to trigger NLRP3 inflammasome activation in a KCNN4-dependent manner. Furthermore, KCNN4 inhibition alleviates auto-inflammation in a CAPS-mimicking mouse model. In other NLRP3-dependent inflammatory conditions that are primarily caused by specific NLRP3 activators that may or may not trigger K^+^ efflux, KCNN4 activation may have rather synergistic effects on NLRP3 inflammasome activation. In this study, we showed that MSU-induced arthritis in mice is attenuated in the absence of PIEZO-dependent mechanotransduction, clearly demonstrating the importance of mechanical stress in other NLRP3-dependent inflammatory conditions. Whether inhibition of KCNN4 is indeed beneficial in such a context remains to be explored in the future.

In this study, we found that PIEZO-dependent effects on NLRP3 inflammasome activation depended on extracellular Ca^2+^. The role of Ca^2+^ in the activation cascade of the NLRP3 inflammasome in response to several NLRP3 activators including nigericin and ATP is controversially discussed (*43*). Importantly however, we emphasize here that Ca^2+^ influx or PIEZO activation is not required per se for NLRP3 inflammasome activation in response to such stimuli when used at higher concentrations. Yet, we provide evidence for a mechanism in which enhancing K^+^ efflux through Ca^2+^-dependent activation of KCNN4 lowers the threshold of NLRP3 inflammasome activation. Activation of PIEZO1 is therefore efficient at concentrations of nigericin that are not sufficient for robust NLRP3 inflammasome activation on its own. Our data thus do not contradict a generally accepted model in which K^+^ is the primary trigger of NLRP3 inflammasome activation. Most importantly, we demonstrate that our findings are of high relevance in the context of NLRP3-dependent inflammatory disorders.

So far, mechanisms underlying Ca^2+^-dependent NLRP3 activation remained ill-defined. One study suggested that Ca^2+^ promotes interaction of ASC with NLRP3 (*44*). It was also proposed that elevated Ca^2+^ levels in the cytoplasm resulted in mitochondrial Ca^2+^ overload that impaired mitochondrial function, thereby activating the NLRP3 inflammasome (*45*). We propose here that Ca^2+^ primarily amplifies K^+^ efflux in a KCNN4-dependent manner, which could be of relevance for other Ca^2+^ mobilizing mechanisms reported during NLRP3 activation.

While our data provide evidence for PIEZO-mediated mechanotransduction to be important in NLRP3 inflammasome activation, a recent study addressed the role of TRPV4, another important mechano-gated ion channel in this context. They find that deletion of *TRPV4* affected crystal-but not non-crystal-induced NLRP3 activation. Mechanistically, they propose that TRPV4 is important for efficient phagocytosis of crystals and phagocytosis-dependent ROS production. Crystal-containing phagosomes are destined for degradation to lysosomes. However, inefficient digestion of crystals in lysosomes results in their accumulation, which causes lysosomal leakage. Deletion or inhibition of TRPV4 limited phagocytosis, thereby attenuating ROS production and lysosomal leakage leading to reduced NLRP3 inflammasome activation (*46*). In line with this study, PIEZO1 activity has also been shown to enhance phagocytosis in macrophages but so far not in the context of crystal uptake and NLRP3 inflammasome activation (*6*). In our study, we put forward a phagocytosis-independent PIEZO function in NLRP3 inflammasome activation. Yet, we cannot exclude that crystal-induced phagocytosis might be also limited in MSU-induced arthritis in mice lacking PIEZO proteins in the myeloid compartment.

In CAPS patients, inflammation is often more prominent in eyes, joints and bones, sites favoring movement-associated mechanical stress. Moreover, urticaria seen in CAPS patients is frequently cold-induced. In this study, we discovered that PIEZO-mediated signaling is sufficient to activate the NLRP3 inflammasome in cells expressing CAPS-causing NLRP3 mutants and patient CAPS cells. Furthermore, we showed that KCNN4 inhibition can efficiently alleviate auto-inflammation in a tamoxifen-induced CAPS-mimicking mouse model, which is a model for human CAPS caused by somatic *NLRP3* mutations (*47*). Gouty arthritis is yet another condition where mechanical impact may play an important role. Indeed, we demonstrated that genetic ablation of PIEZO function in mice reduces MSU-induced joint inflammation to a level seen for NLRP3 deletion. Thus, targeting of this pathway may provide a novel therapeutic strategy for treatment of CAPS patients and potentially other NLRP3-related inflammatory diseases.

## Materials and Methods

### Mice

*Piezo1^fl/fl^* (Strain #:029213), *Piezo2^fl/fl^* (Strain #:027720) and *Nlrp3^A350VneoR^* (Strain #:017969) mice were obtained from The Jackson Laboratory. *Piezo1^fl/fl^* and *Piezo2^fl/fl^*mice were crossed with *LysM-Cre* mice to obtain mice with myeloid-specific deletion of *Piezo1* and *Piezo2*. *Cx3cr1-CreER* mice were from Prof. Marco Prinz at University of Freiburg, Germany. *Nlrp3^A350VneoR^* mice were crossed with *Cx3cr1-CreER* mice to obtain *Nlrp3^A350VneoR^*;*Cx3cr1-CreER* mice with tamoxifen-inducible expression of NLRP3^A350V^ in Cx3cr1-positive cells. Mice were housed under specific pathogen-free conditions with controlled temperature (19-23°C) and humidity (50-60%) on a 12-h light/dark cycle with unrestricted access to water and standard laboratory chow. Maintenance and animal experimentation were in accordance with the local ethical committee (Com’Eth) in compliance with the European legislation on care and use of laboratory animals (La cellule AFiS (Animaux utilisés à des Fins Scientifiques): APAFIS#36729-2022041911158105 v2 and APAFIS #43633-2023042814302902 v6). No exclusion of animals used for experiments was performed. Littermates were chosen randomly according to their genotypes.

### Reagents

Imiquimod (R837) (tlrl-imq), MSU crystals (tlrl-msu-25), CL097 (tlrl-c97) and MCC950 (inh-mcc) were purchased from InvivoGen. Nigericin sodium salt (N7143), Propidium Iodide (P4170), Lipopolysaccharides from Escherichia coli 055: B5 (L2880) and Phorbol 12-myristate 13-acetate (PMA) (P1585) were obtained from Sigma-Aldrich. Yoda1 (5586) and TRAM34 (2946) were purchased from Bio-techne. Sytox™ Green Nucleic Acid Stain (S7020) was purchased from Thermo Fisher Scientific. Human macrophage-colony stimulating factor (hM-CSF) (11343117) was obtained from Immunotools. Anti-Caspase1 (p20) (human) antibody (AG-20B-0048-C100), anti-Caspase1 (p20) (mouse) antibody (AG-20B-0042), anti-NLRP3 antibody (AG-20B-0014-C100) were purchased from AdipoGen. Anti-human IL-1β antibody (AF-201-NA) was from R&D Systems. Anti-murine IL-1β antibody (5129-100) was from BioVision. Anti-tubulin antibody (T5168) was from Sigma-Aldrich. Anti-GFP antibody (ab13970) and anti-NEK7 antibody (ab133514) were obtained from Abcam. Anti-ASC antibody (sc-22514) was obtained from Santa Cruz Biotechnology. Anti-Gasdermin D antibody (NBP2-33422) was obtained from Novus Biologicals.

### Plasmids

The pX330-P2A-EGFP and pX330-P2A-RFP plasmid was previously generated by inserting P2A-GFP and P2A-RFP sequence into pX330-U6-Chimeric_BB-CBh-hSpCas9 (*26*). KCNN4 CDS was amplified from THP-1 cDNA by PCR. KCNN4 fused with EGFP was cloned into pBOB plasmid using ligation-independent cloning (LIC). Calcium reporter jGCaMP7s sequence was amplified from pGP-CMV-jGCaMP7s (104463, Addgene) and cloned into pBOB plasmid by LIC. C-terminal EGFP-tagged CAPS-causing mouse Nlrp3 mutants R258W, D301N and T346M were cloned into pBOB plasmid by LIC. Mouse Piezo1 CDS was amplified from a gift plasmid from Prof. Ardem Patapoutian and cloned into pBOB plasmid by LIC. The primers used are listed below:

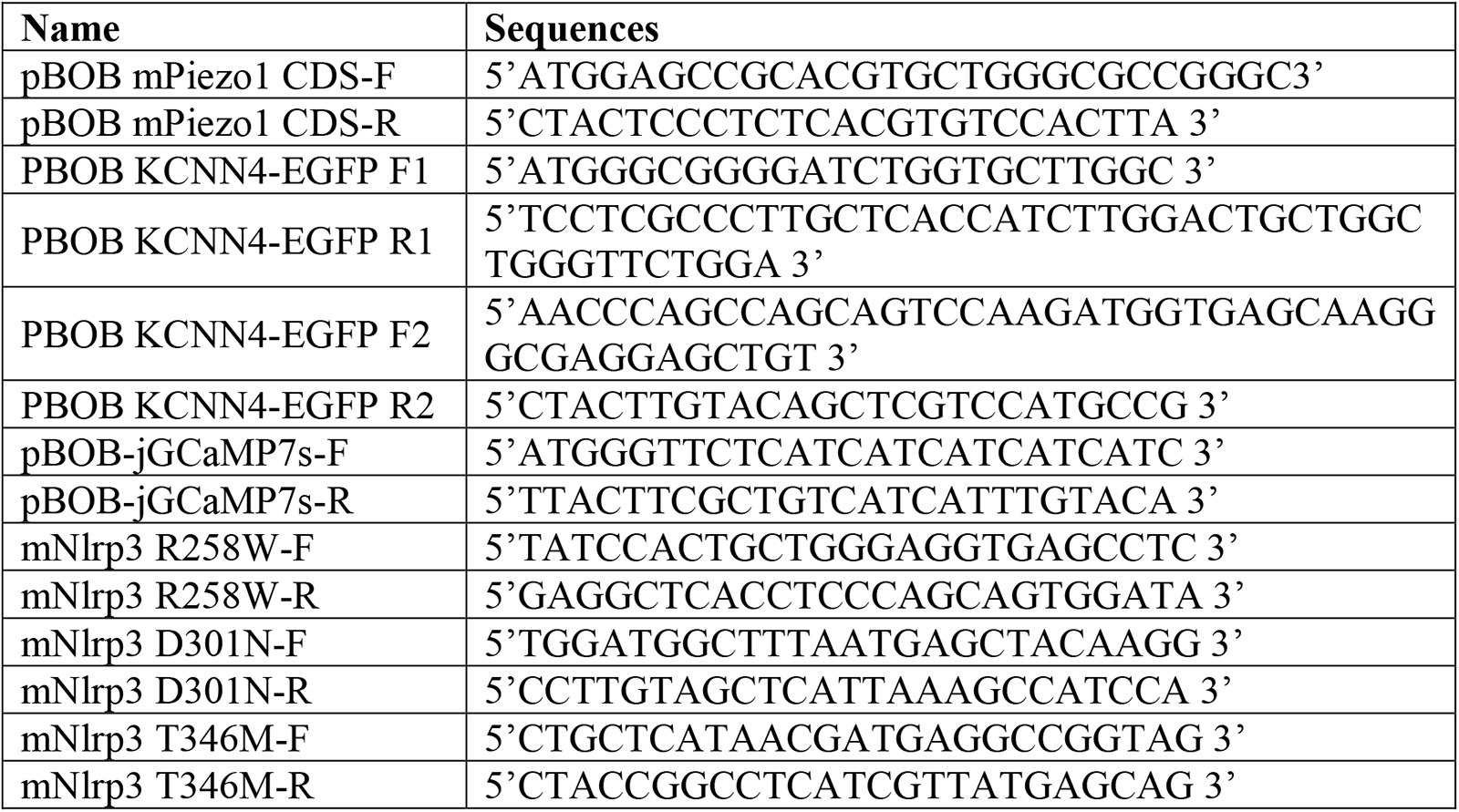

### Generation of knockout cell lines using CRISPR/Cas9-mediated gene editing

For the generation of *PIEZO1* KO, *PIEZO2* KO, *PIEZO1/2* double knockout (dKO) and *KCNN4* KO THP-1 cell lines, two sgRNAs (sgRNA 1 and sgRNA 2) for each gene were designed and cloned into pX330-P2A-EGFP or pX330-P2A-RFP, respectively, using T4 ligation. THP-1 cells were transfected with mixture of two sgRNAs-expressing plasmids (0.5 µg each) using X- tremeGENE 9 according to the manufacturer’s manual. 24 h after transfection, GFP and RFP double-positive cells were sorted and collected using BD FACS Aria™ III Cell Sorter. Single cell colonies were obtained by seeding into 96-well plates via series of dilution. The sequences of sgRNAs are listed below:

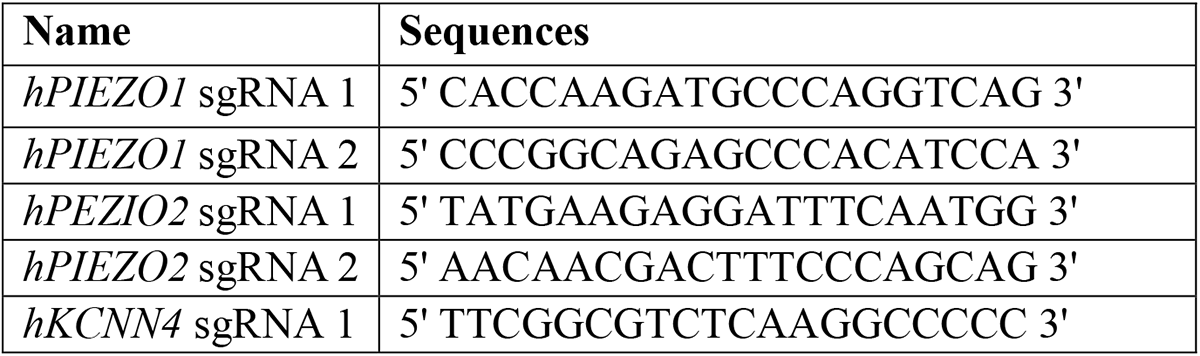
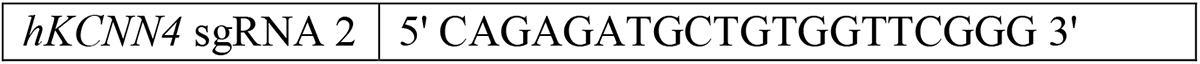

### Validation of obtained THP-1 knockout cell lines by Sanger sequencing

For the validation of *PIEZO1* KO, *PIEZO2* KO, *PIEZO1/2* dKO, *KCNN4* KO THP-1 clones, genomic fragments containing sgRNA targeting sites were amplified by PCR and cloned into pUC57 plasmid using LIC. After transformation of *E. coli* competent cells, plasmids were purified from at least 6 bacteria colonies and subjected to Sanger sequencing. Obtained sequences were aligned to reference sequence to determine the insertion/deletion. The primers used are listed below:

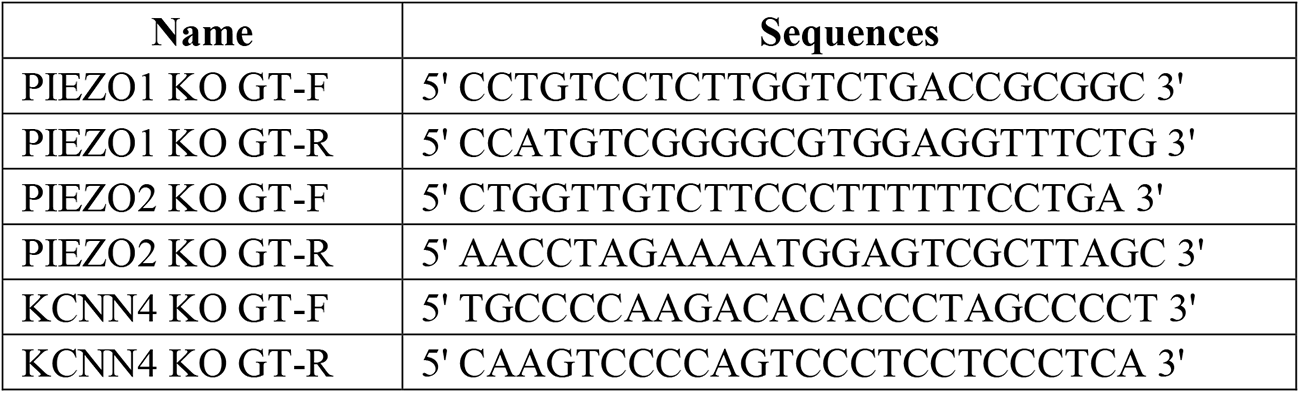

### Cell culture

THP-1, mouse primary bone marrow-derived macrophages (BMDMs) and HEK293t cell lines were cultured at 37°C with 5% CO2. THP-1 cells were grown in RPMI 1640 supplement with 10 mM HEPES, 10% fetal calf serum (FCS), 2.5 g/l glucose, 1 mM sodium pyruvate, gentamycin and 50 µM β-mercaptoethanol. HEK293t cells were grown in DMEM (1 g/ml glucose) supplement with 10% fetal calf serum, penicillin and streptomycin. BMDMs were differentiated from bone marrow progenitors isolated from the tibia and femur in DMEM (4,5g/l glucose) supplement with 50 ng/ml recombinant hM-CSF, 10% heat-inactivated fetal calf serum, penicillin and streptomycin for 7 days. Human bloods from heathy control and CAPS patient were collected after the patient gave their signed informed consent. Peripheral blood mononuclear cells (PBMCs) were isolated using Ficoll-Paque PLUS (GE Healthcare) and cultured in RPMI 1640 containing 10% heat-inactivated fetal bovine serum, penicillin and streptomycin. For treatment of cells, BMDMs and THP-1 cells were primed with 1 µg/mL LPS for 3 h, followed by the treatment of nigericin, Imiquimod and CL097 in presence or absence of 25 µM Yoda1. For treatment of inhibitor, cells were pretreated with inhibitor for 45 min and followed by stimulation in presence of inhibitor. To assess the involvement of K^+^ efflux, LPS-primed cells were treated with NLRP3 inflammasome activator in the culture medium supplemented with 5 mM, 15 mM, 25 mM, or 35 mM KCl. For the experiments with calcium-free medium, BMDMs and THP-1 cells were primed with 1 µg/mL LPS for 3h in DMEM (4,5g/l glucose) containing 10% FCS, cells were washed once with PBS before treatment with nigericin, R837 or CL097 in DMEM (4,5g/l glucose) without calcium containing 10% dialyzed FCS.

For the stiffness-related experiments using wild type THP-1 cells, cells were seeded on silicone gels of 0.2 KPa and 64 KPa (5165-5EA and 5145-5EA, Sigma-Aldrich) with culture medium containing 100 nM PMA for 3 h, followed by replacement of fresh medium and overnight incubation at 37°C. On the next day, cells were treated with nigericin or R837. For the stiffness-related experiments using THP-1 cells expressing CAPS-causing Nlrp3 mutants, cells were seeded on normal tissue culture plates, 0.2 KPa or 64 KPa silicone gels with culture medium containing 100 nM PMA for 3 h, followed by replacement of fresh medium and supernatants were collected after 4 hours incubation at 37°C.

For shear stress related experiments, shear stress was generated by a rotating unit device as described previously (*48*). Briefly, each rotating unit is made of one circular rotating plate in Plexiglas (3.2 cm diameter) fixed to a rotating motor. Rotating plate is replaced in contact with the culture medium which generates a shear flow over cells when rotates. The shear stress *τ* exposed to cells depends on the circular plate rotation speed *Ω*, the distance from the axis *r* and the height of the plate *h* and is given by: *τ = η Ω r/h*, *η* being the medium viscosity. In these experiments, the rotating plates were set up at height *h* = 4 mm and at speed *Ω* = 8 and 16 rounds *per* min, which generates maximal shear stress at the edge of rotating plate τ *≈ 1.8 × 10^-3^ Pa* and *3.6 × 10^-3^ Pa*, respectively. WT and *PIEZO1/2* dKO cells were seeded on 6-well tissue culture plates with culture medium containing 100 nM PMA for 3h, followed by replacement of fresh medium and overnight incubation at 37°C. On the next day, cells were treated with nigericin or R837 in presence or absence of indicated shear stress.

### Immunoblotting

Both cell lysates and culture supernatants were subjected to immunoblotting. For cell lysates, cells were lysed on ice with 1× RIPA buffer (50 mM Tris-HCl pH 7.5, 150 mM NaCl, 1 mM EDTA, 1 mM EGTA, 1% Triton X-100, 1 mM NaVO4, 1.5 mM Sodium pyrophosphate, 1 mM NaF, 1 mM β-Glycerophosphate) supplemented with protease inhibitor cocktail. The immunoblot was prepared using Glycine SDS-PAGE gels. For the supernatants, the proteins were extracted using methanol-chloroform precipitation protocol as previously described (*26*), separated by Tricine SDS-PAGE and analyzed by immunoblotting. The PVDF membranes were incubated with primary antibody at 4 °C overnight. After 3 times washes with TBS-T, membranes were incubated with HRP-conjugated secondary antibody for 1 hour at RT. Membranes were again washed with TBS-T for 3 times and were incubated with Immobilon Forte Western HRP substrate (WBLUF0500, EMD Millipore Corporation). Images were captured using AI600 Imager from GE Healthcare Life Science.

### Flow cytometry

For analysis of lytic cell death, after treatments, THP-1 cells were spin down and resuspended in cold PBS with 1 µM of Sytox Green or propidium iodide (PI). Sytox Green-or PI-positive cells were analyzed by BD FACS Celesta™ Flow Cytometer. For cell sorting, THP-1 cells were spin down and resuspended in cold PBS supplemented with 1% of fetal calf serum, followed by sorting at sterile condition using BD FACS Aria™ III Cell Sorter in fresh medium for further culture.

### Measurement cytokines using ELISA

IL-1β and TNFα levels in collected culture supernatants were determined by ELISA according to the user’s manual. Human IL-1 beta/IL-1F2 DuoSet ELISA (DY201), mouse IL-1 beta/IL-1F2 DuoSet ELISA (DY401), human TNF-alpha DuoSet ELISA (DY210), mouse TNF-alpha DuoSet ELISA (DY410) and DuoSet ELISA Ancillary Reagent Kit 2 (DY008) were purchased from R&D system.

### Immunofluorescence and live-video imaging

For immunofluorescence, cells plated on coverslips (12-mm) were fixed with 4% paraformaldehyde for 15 min at room temperature after treatments. Coverslips were washed three times with PBS for 5 min each time and permeabilized with 0.1% saponin in PBS for 10 min, followed by blocking in PBST (PBS plus 0.05% Tween-20) containing 0.5% BSA for 1 h. The coverslips were further incubated with anti-ASC antibody (1/100 dilution) for 1 h at room temperature, followed by incubation with the secondary for 1h at room temperature (avoid the coverslips in direct light). After three times washing with PBST, cells were stained with DAPI and mounted for further imaging. For live-video imaging, THP-1 WT and *PIEZO1/2* dKO cells stably expressing jGCaMP7S were seeded on µ-dish 35 mm high glass bottom (81158, Ibidi) in culture medium containing 100 nM PMA for 3 h, followed by replacement of fresh medium and incubation overnight. On the next day, cells were treated with R837, Yoda1 or R837 plus Yoda1. The images were acquired using a Nikon spinning disk with an interval of 20s. Ionomycin was added at ∼16 min after treatment. Fluorescence intensities of individual cells over time were quantified with Image J.

### Lentivirus packaging and infection

The 3^rd^ generation lentiviral vector system with three packaging helper plasmids (pVSVG, pMDL and pREV) was used in this study. The lentivirus packaging was performed in HEK293t cells. HEK293t cells were transfected with Lipofectamine 2000 according to the user’s manual. THP-1 cells (4.0 × 10^5^ cells per well in 6-well plate) were infected in 1.5 mL lentivirus-contained culture medium plus 1 mL fresh medium containing final concentration of 10 µg/mL polybrene. 48 hours after viral infection, cells were used for further experiments.

### Genome-wide CRISPR/Cas9-mediated knockout screen

The Brunello human genome-wide CRISPR/Cas9 knockout lentiviral library containing 76,441 sgRNAs systemically targeting 19,114 genes on human genome was used in this study. Briefly, THP-1 cells stably expressing Cas9 protein were infected with the pooled Brunello lentivirus library at ROI of 0.2. After puromycin selection for 3 days, cells were further expanded. To ensure a representation of minimal 500 cells per sgRNA, at least 4.0×10^7^ cells were used in each group. An adequate number of cells were primed with LPS in suspension for 3 hours and treated with vehicle (control group) or R837 plus Yoda1 (treated group). When pyroptotic cell death reached ∼85% in the treated group, R837 and Yoda1 were washed out and the survivals were expanded. When cell number reached 4.0 × 10^7^ cells, these cells were repeatedly treated with R837 plus Yoda1 to reach reached ∼85% of pyroptotic cell death. R837 and Yoda1 were washed out and the survivals were further expanded to reach 4.0 × 10^7^ cells. Genomic DNA from both control- and treated groups was isolated using the phenol/chloroform extraction method. Fragments containing sgRNA were amplified from genomic DNA by PCR referring to the protocol from broad institute GPP Web Protal (https://portals.broadinstitute.org/gpp/public/resources/protocols). PCR fragments were cleaned up using NucleoSpin gel and PCR clean-up kit (740609.50, MACHEREY-NAGEL) according to the users’ manual and subjected to next generation sequencing (NGS). Sequencing was performed on an Illumina HiSeq 4000 sequencer as 50 bp single end reads by the GenomEast platform, a member of the ‘France Genomique’ consortium (ANR-10-INBS-0009). Image analysis and base calling were performed using RTA version 2.7.7 and bcl2fastq version 2.20.0.422. Sequencing reads were trimmed, aligned and counted using PinAPL-Py and guide RNAs (Brunello library). The Matching Threshold for bowtie2 alignment was defined as 35. Data were statistically analyzed using Model-based Analysis of Genome-wide CRISPR/Cas9 Knockout (MAGeCK v0.5.9.3). The method ranks sgRNAs based on *p*-values calculated from the negative binomial model, and uses a modified robust ranking aggregation (RRA) algorithm to identify positively or negatively selected genes. Read counts were normalized by total number of reads. Enrichment testing of paired samples was performed using the test function in MAGeCK with -- paired option. Volcano plot was generated using ggplot package in R and significance was defined as FDR < 0.05.

### MSU-induced acute arthritis mouse model

MSU crystal (0.5 mg in 10 µl PBS) was injected intra-articularly into one ankle of each adult mouse. The thickness of injected ankles was monitored using a caliper at 0, 2, 6, 12 and 24 hours after MSU injection. Ankle temperature was measured using an infrared thermometer at 12 hours after MSU injection.

### CAPS-mimicking mouse model with nonlethal autoinflammation

Adult mice were daily treated with tamoxifen 50 mg/kg via intraperitoneal injection for five consecutive days. Then, mice were further daily treated with vehicle (10% DMSO:90% corn oil) or TRAM34 dissolved in 10% DMSO:90% corn oil (15 mg/kg) via intraperitoneal injection for eight days. Body weight loss was monitored every day starting from the first day of TRAM34 treatment. Rectal temperature was measured and blood was collected on the day after the last day of TRAM34 treatment. The timeline of this experimental procedure is illustrated in Fig.7A.

### Statistical analysis

Preliminary experiments were performed and sample size was determined based on generally accepted rules to test preliminary conclusions reaching statistical significance, where applicable. For FACS experiments, the percentage of Sytox green-positive and PI-positive cells was provided. For immunofluorescence experiments, a minimum of 150 cells were counted per condition, and the percentage containing ASC specks cells was calculated. A non-parametric *Mann-Whitney* test (two-tailed) was used for statistical analysis of ankle temperature, rectal temperature, and serum IL-1β levels. Differences between groups were assessed for statistical significance by ANOVA or using an unpaired Student *t* test (when only two sets of data were compared). Statistical significance was indicated using the following symbols: * *p* < 0.05, ** *p* < 0.01 and *** *p* < 0.001.

## Supplementary Materials

**Fig. S1.**
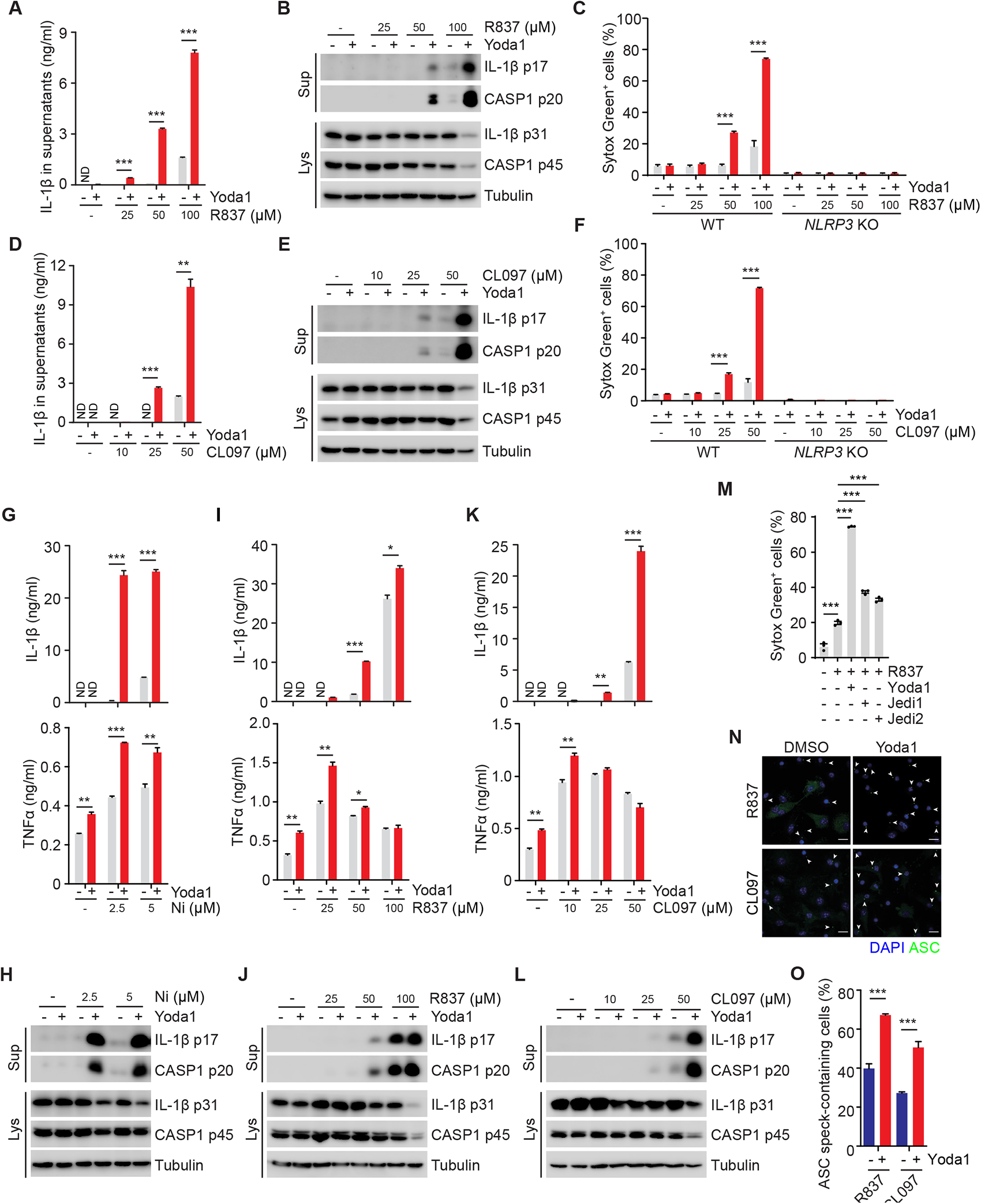
Activation of PIEZO1 by Yoda1 potentiates R837-and CL097-induced NLRP3 inflammasome activation. (**A**, **D**) ELISA measurements of IL-1β in culture supernatants from THP-1 cells primed with 1 µg/ml LPS for 3 h and followed by treatment with R837 (**A**) or CL097 (**D**). (**B**, **E**) Immunoblotting of culture supernatants (Sup) and lysates (Lys) from LPS-primed THP-1 treated as described for panel A and D. Antibodies againist IL-1β and Caspase-1 (CASP1) were used. Antibody against Tubulin was used as a loading control. (**C**, **F**) Uptake of Sytox Green in LPS-primed WT and *NLRP3* KO THP-1 cells treated as described for panel A and D. (**G**, **I**, **K**) ELISA measurements of IL-1β and TNFα in culture supernatants from LPS-primed BMDMs treated with nigericin (Ni) (**G**), R837 (**I**), CL097 (**K**) as indicated in presence or absence of 25 µM Yoda1. (**H**, **J**, **L**) Immunoblotting of culture supernatants (Sup) and lysates (Lys) from LPS-primed BMDMs treated as in panel G, I, K. Antibodies againist IL-1β and Caspase-1 (CASP1) were used. Antibody against Tubulin was used as a loading control. (**M**) Uptake of Sytox Green in LPS-primed WT THP-1 cells treatment with 100 µM R837 in presence or absence of 25 µM Yoda1, 100 µM Jedi1 or 100 µM Jedi2. (**N**) Representative immuno-fluorescence images of ASC speck formation in LPS-primed BMDMs stimulated with 100 μM R837 or 50 μM CL097 in the presence or absence of 25 µM Yoda1. White arrows indicate ASC specks. Scale bars: 10 µm. (**O**) Quantification of macrophages containing ASC specks in panel N. “ND” not detected. \**p < 0.05*, \*\**p < 0.01*, \*\*\**p < 0.001*. Data are representative of at least three independent experiments.

**Fig. S2.**
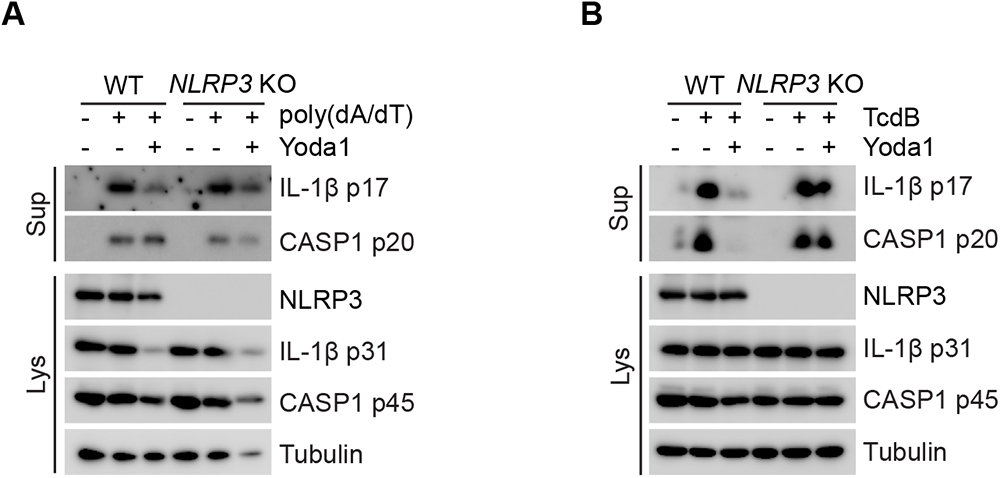
Yoda1 does not enhance the activity of AIM2- and Pyrin inflammasome. (**A**, **B**) Immunoblotting of culture supernatants (Sup) and lysates (Lys) from LPS-primed WT and *NLRP3* KO BMDMs transfected with 1µg/ml poly(dA:dT) for 4 h (**A**) or treated with 1 nM TcdB for 2 h (**B**) in presence or absence of 25 µM Yoda1. Antibodies againist IL-1β and Caspase-1 (CASP1) were used. Antibody against Tubulin was used as a loading control. Data are representative of three independent experiments.

**Fig. S3.**
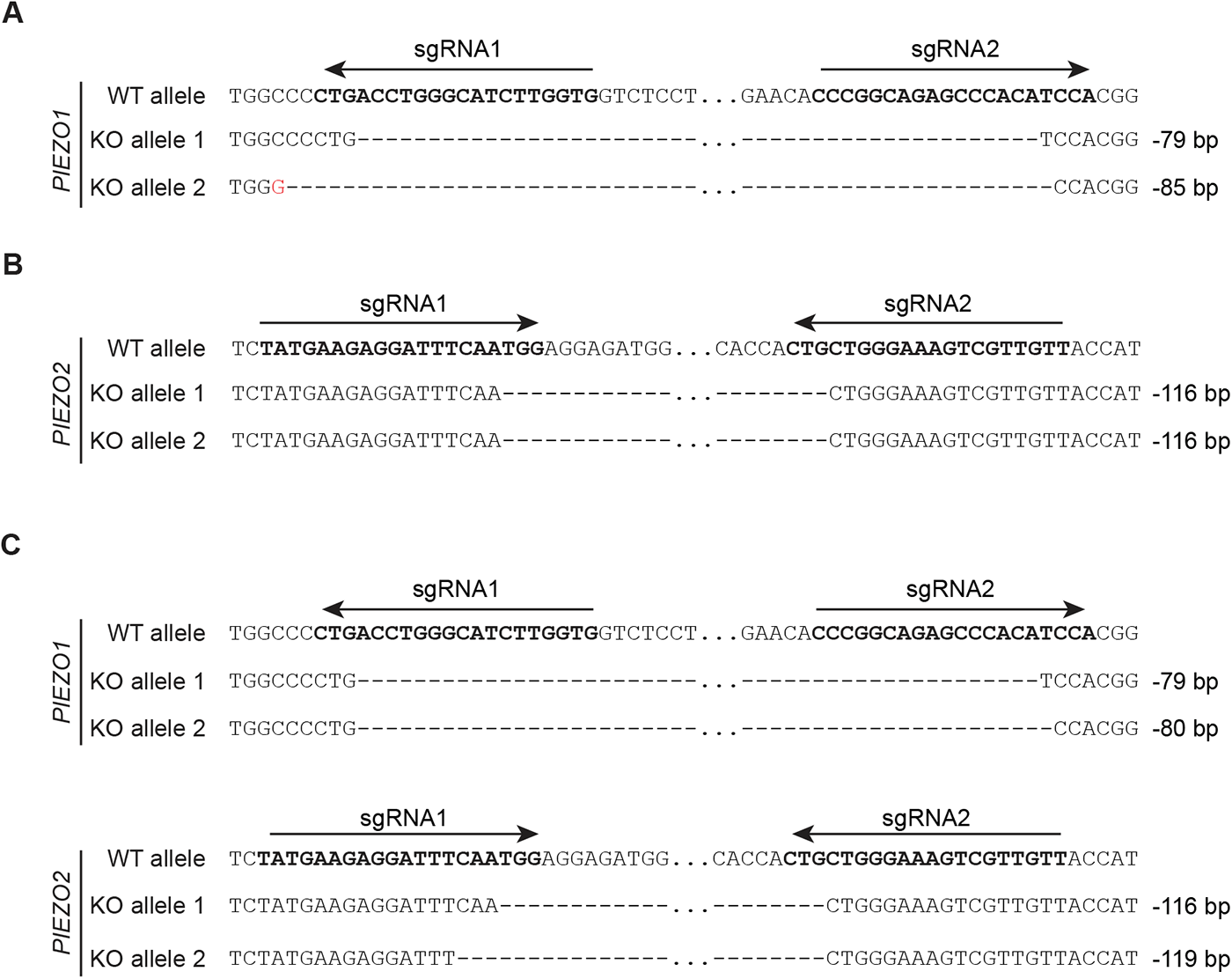
Validation of *PIEZO1* KO, *PIEZO2* KO and *PIEZO1/2* dKO THP-1 cells. (**A**-**C**) Sequences of *PIEZO1* and/or *PIEZO2* alleles validated by Sanger sequencing in *PIEZO1* KO (**A**), *PIEZO2* KO (**B**), *PIEZO1/2* dKO (**C**). The sequences of sgRNAs are indicated by arrows. A mutation is highlighted in red, “-” indicates a deleted nucleo-tide, “…” in *PIEZO1* alleles indicates a fragment of 34 bps; “…” in *PIEZO2* alleles indicates a fragment of 96 bps. The seqeuncing was performed once.

**Fig. S4.**
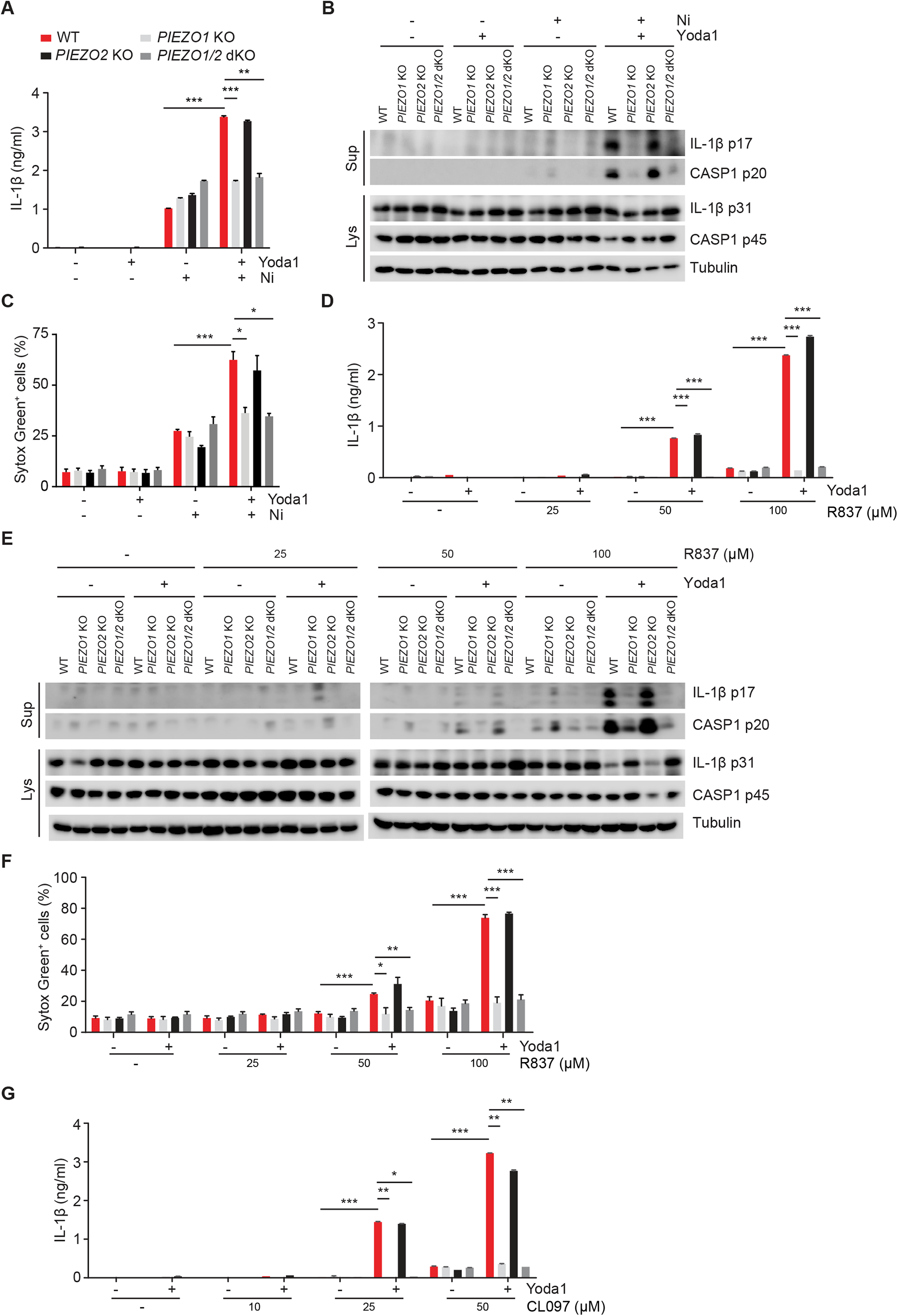
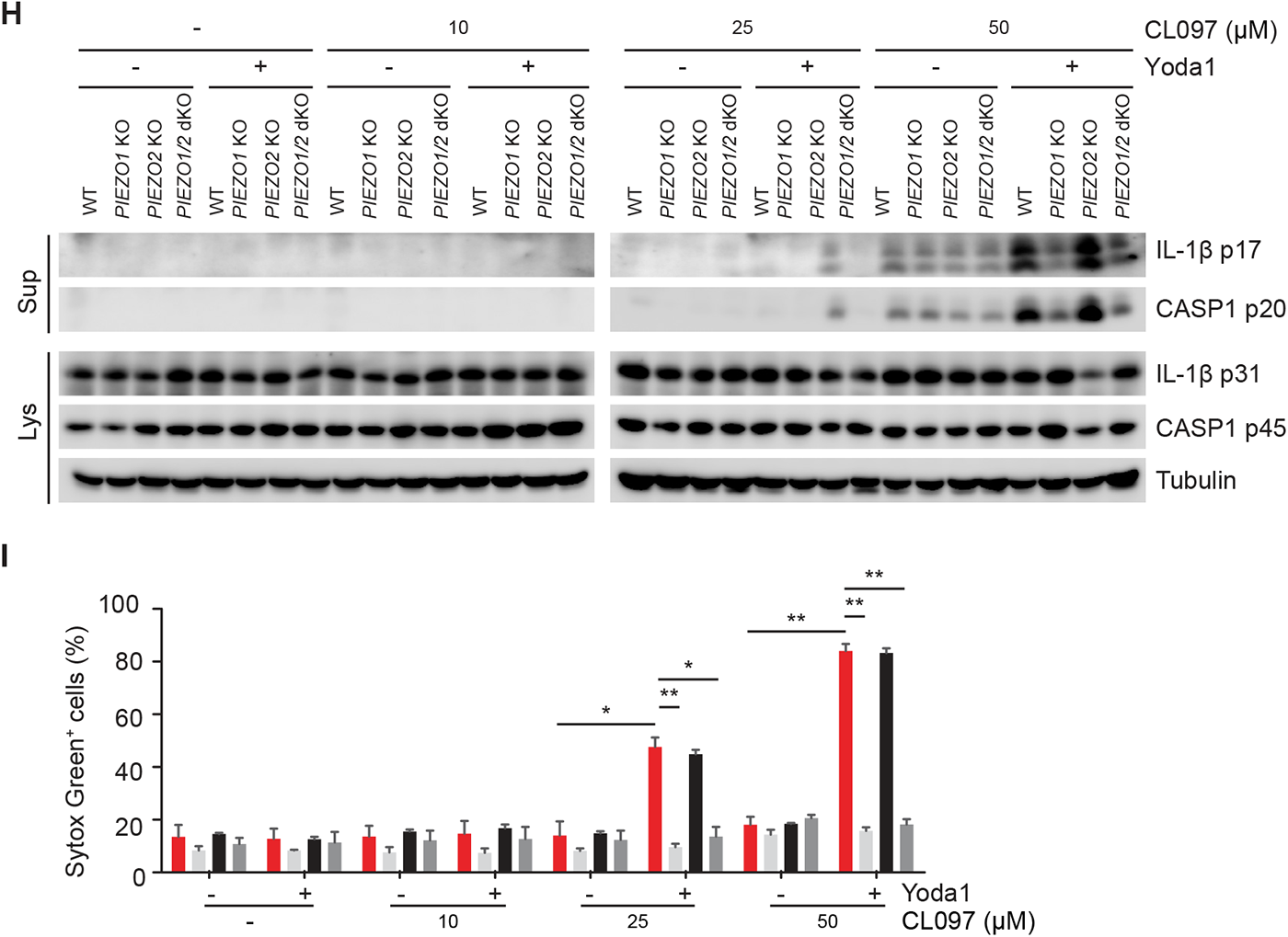
Deletion of *PIEZO1* abolishes Yoda1-dependent NLRP3 inflammasome activation. (**A**, **D**, **G**) ELISA measurements of IL-1β in culture supernatants from WT, *PIEZO1* KO, *PIEZO2* KO and *PIEZO1/2* dKO THP-1 cells primed with 1 µg/ml LPS for 3 h and followed by treatment with nigericin (**A**), R837 (**D**) and CL097 (**G**) as indicated in presence or absence of 25 µM Yoda1. (**B**, **E**, **H**) Immunoblotting of culture supernatants (Sup) and lysates (Lys) from LPS-primed THP-1 cells in experiments as described for panel A, D and G. Antibodies againist IL-1β and Caspase-1 (CASP1) were used. An antibody against Tubulin was used as a loading control. (**C**, **F**, **I**) Uptake of Sytox Green from LPS-primed cells in experiments as described in panel A, D and G. Sytox Green uptake was analyzed by FACS after staining. \**p < 0.05*, \*\**p < 0.01*, \*\*\**p < 0.001*. Data are representative of at least three independent experiments.

**Fig. S5.**
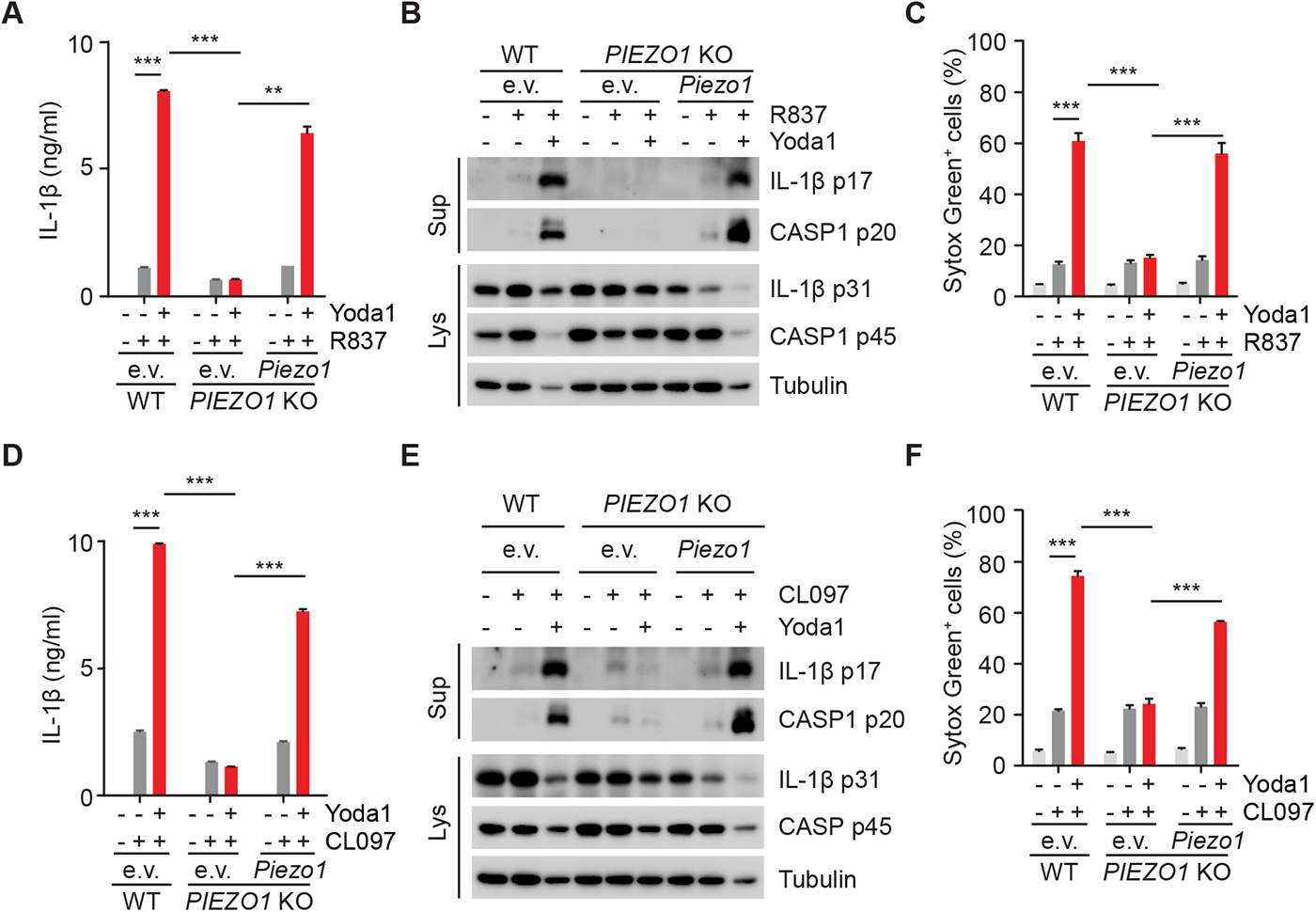
Re-expression of Piezo1 restores Yoda1-dependent NLRP3 inflammasome activation in *PIEZO1* KO cells. (**A**, **D**) ELISA measurements of IL-1β in culture supernatants in WT and *PIEZO1* KO THP-1 cells expressing empty vector (e.v.) or mouse Piezo1. Cells were primed with 1 µg/ml LPS for 3 h, followed by treatment with 100 µM R837 (**A**), 50 µM CL097 (**D**) in presence or absence of 25 µM Yoda1. (**B**,**E**) Immunoblotting of culture supernatants (Sup) and lysates (Lys) from LPS-primed THP-1 cells in experiments as described in panel A and D. Antibodies againist IL-1β and Caspase-1 (CASP1) were used. Antibody against Tubulin was used as a loading control. (**C**, **F**) Uptake of Sytox Green from LPS-primed cells in experiments as described for panel A and D. Sytox Green uptake was analyzed by FACS after staining. \*\**p < 0.01*, \*\*\**p < 0.001*. Data are representative of at least three independent experiments.

**Fig. S6.**
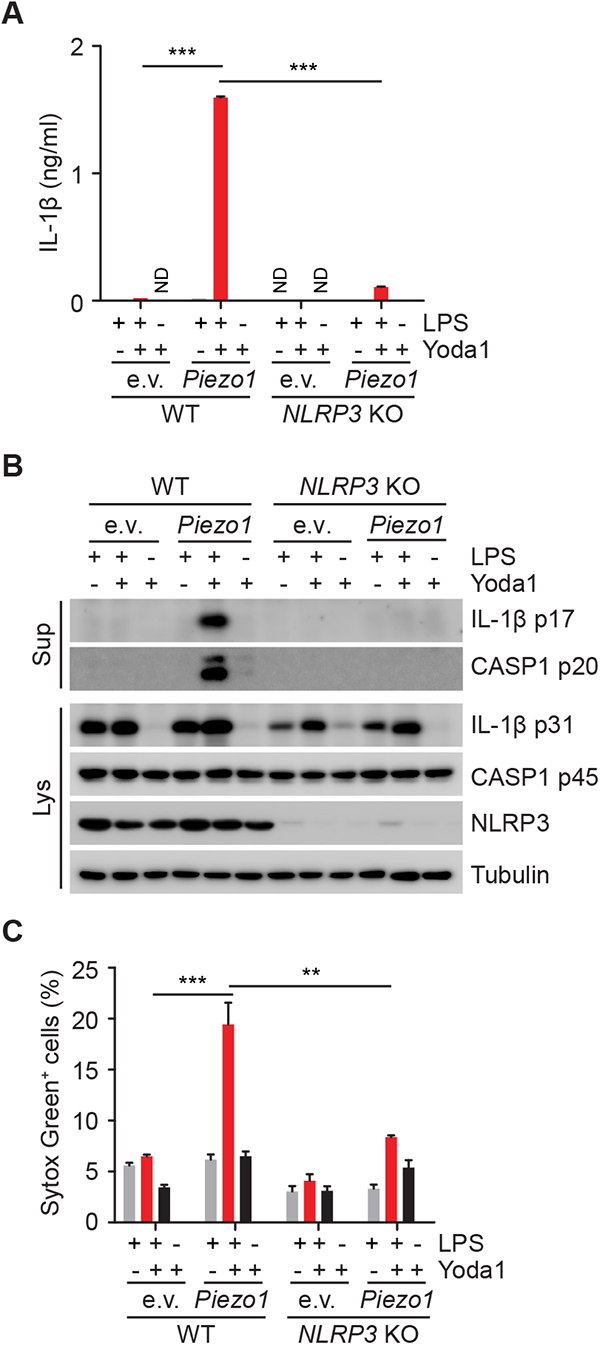
Yoda1 is sufficient to activate NLRP3 inflammasome in cells ectopically-expressing Piezo1. (**A**) ELISA measurements of IL-1β in culture supernatants in WT and *NLRP3* KO THP-1 cells expressing empty vector (e.v.) or mouse Piezo1. Cells were primed with or without 1 µg/ml LPS for 3 h, followed by treatment with 25 μM Yoda1. (**B**) Immunoblotting of culture supernatants (Sup) and lysates (Lys) from cells in experiments as described in panel A. Antibodies againist IL-1β and Caspase-1 (CASP1) were used. An antibody against Tubulin was used as a loading control. (**C**) Uptake of Sytox Green in cells in experiments as described in panel A. Sytox Green uptake was analyzed by FACS after staining. “ND” not detected. \*\**p < 0.01*, \*\*\**p < 0.001*. Data are representative of at least three independent experiments.

**Fig. S7.**
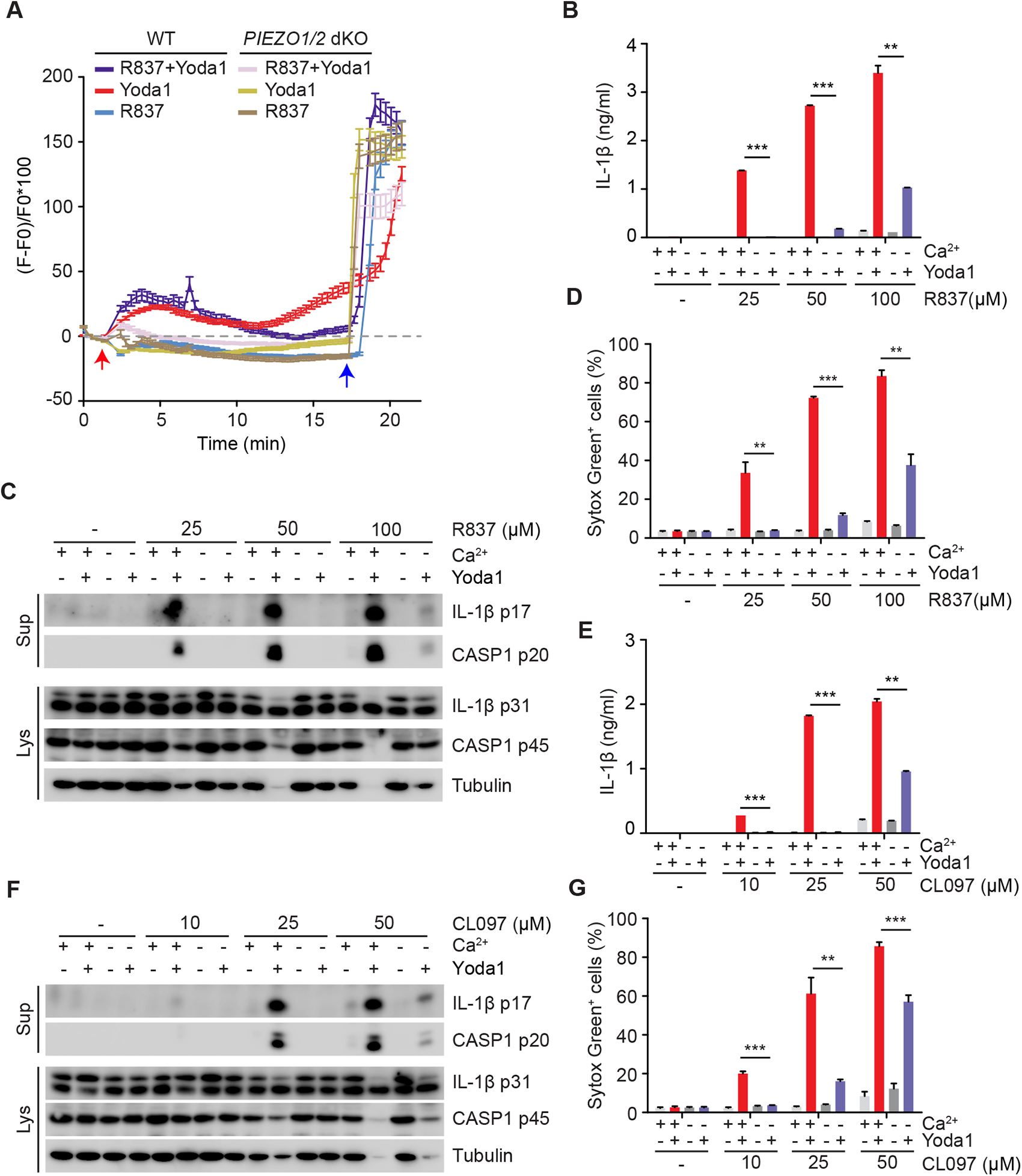
Ca^2+^ influx is required for PIEZO-dependent NLRP3 inflammasome activation. (**A**) Measurement of cytosolic Ca^2+^ levels in WT and *PIEZO1/2* dKO THP-1 cells using the Ca^2+^ reporter jGCaMP7s. Cells were treated with 100 µM R837, 25 µM Yoda1 or 100 µM R837 plus 25 µM Yoda1. The images were acquired using a Nikon spinning-disk microscope with an interval of 20s. Stimuli were added into the culture medium at 1 min as indicated by a red arrow, and ionomycin was added at the end of the experiments as indicated by a blue arrow. Fluorescence intensities of individual cells over time were quantified. n=79 for “WT R837+Yoda1” group; n=96 for “WT Yoda1” group; n=90 for “WT R837” group; n=113 for “*PIEZO1/2* dKO R837+Yoda1” group; n=83 for “*PIEZO1/2* dKO Yoda1” and n=91 for “*PIEZO1/2* dKO R837” group. (**B**, **E**) ELISA measurements of IL-1β in culture supernatants from LPS-primed THP-1 treated with R837 (**B**), CL097 (**E**) as indicated in presence or absence of 25 µM Yoda1 in medium with Ca^2+^ or without (w/o) Ca^2+^. (**C**, **F**) Immunoblotting of culture supernatants (Sup) and lysates (Lys) from LPS-primed THP-1 cells in experiments as described for panel B and E. Antibodies against IL-1β and Caspase-1 (CASP1) were used. An antibody against Tubulin was used as a loading control. (**D**, **G**) Uptake of Sytox Green in LPS-primed THP-1 cells in experiments as described in panel B and E. Sytox Green uptake was analyzed by FACS after staining. \*\**p < 0.01*, \*\*\**p < 0.001*. Data are representative of at least three independent experiments.

**Fig. S8.**
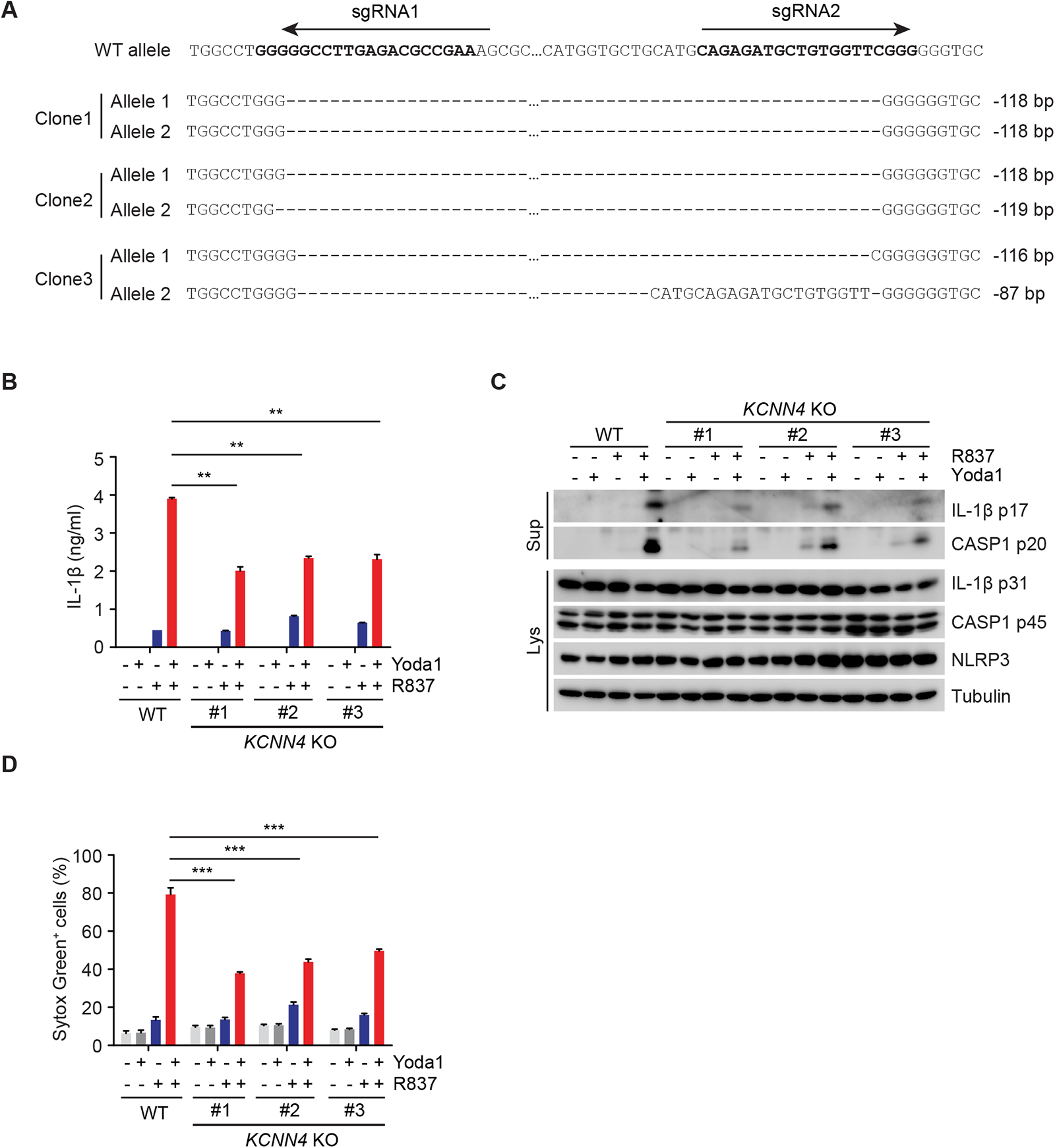
Deletion of KCNN4 abolishes Yoda1-dependent NLRP3 inflammasome activation. (**A**) Generation of THP-1 *KCNN4* KO cells using CRISPR/Cas9-mediated gene editing. Validation of *KCNN4* KO THP-1 cells by Sanger Sequencing. sgRNAs were indicated by arrows. “-” means a deleted nucleotide, “…” means a fragment of 65 bps. (**B**) ELISA measurements of IL-1β in culture supernatants from WT and *KCNN4* KO THP-1 cells primed with 1 µg/ml LPS for 3 h and followed by treatment with 100 µM R837 in presence or absence of 25 µM Yoda1 for 1 h. (**C**) Immunoblotting of culture supernatants (Sup) and lysates (Lys) from LPS-primed THP-1 cells in experiments as described for panel B. Antibodies against IL-1β, Caspase-1 (CASP1) and NLRP3 were used. An antibody against Tubulin was used as a loading control. (**D**) Uptake of Sytox Green in LPS-primed cells in experiments as described for panel B. \*\**p < 0.01*, \*\*\**p < 0.001*. Data are representative of three independent experiments.

**Fig. S9.**
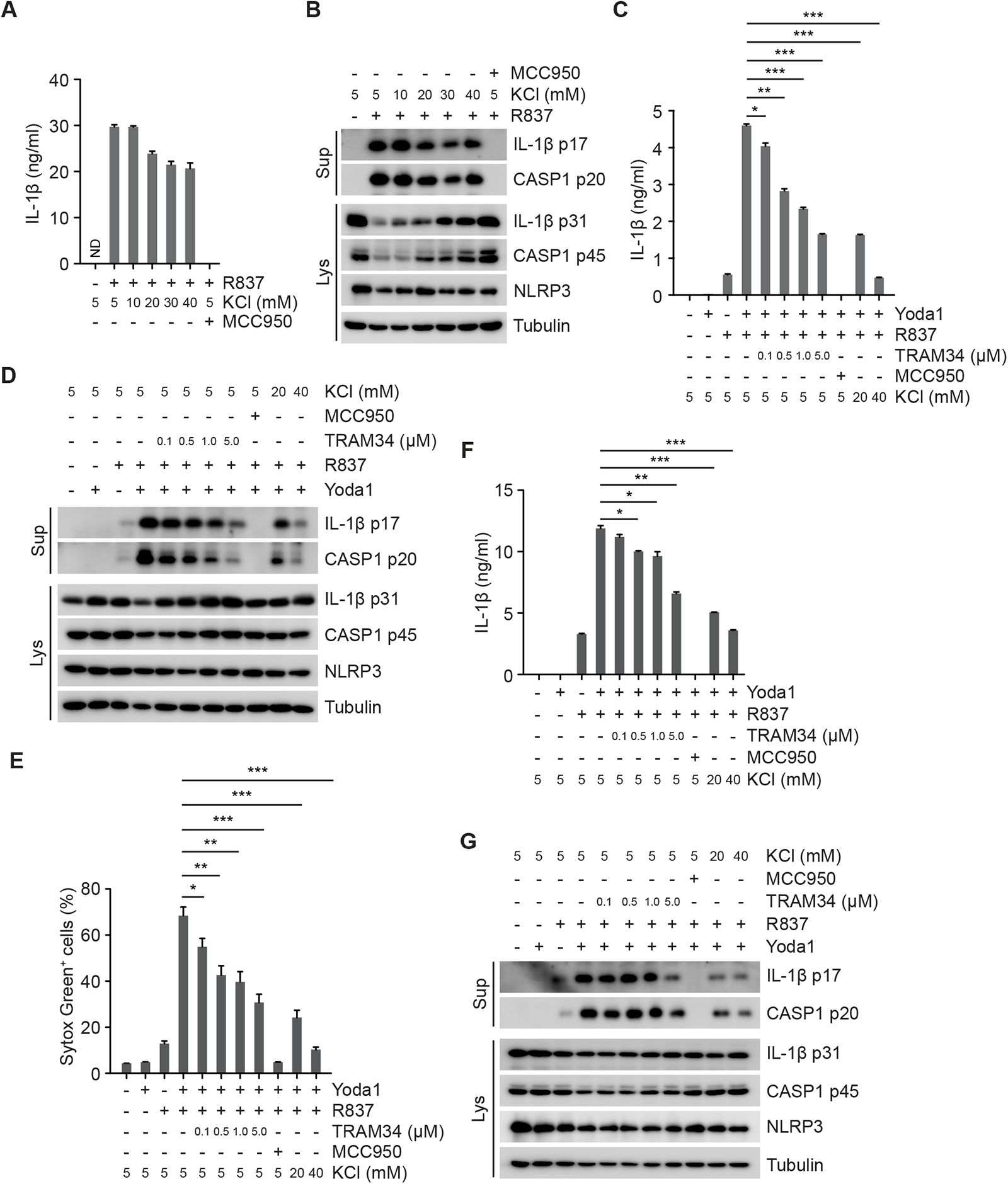
Inhibition of KCNN4 blocks the effect of Yoda1 on NLRP3 inflammasome activation. (**A**) ELISA measurements of IL-1β in culture supernatants from LPS-primed BMDMs treated with 100 µM R837 in medium containing indicated concentrations of extracellular KCl for 3 h. (**B**) Immunoblotting of culture supernatants (Sup) and lysates (Lys) from LPS-primed BMDMs in experiments as described for panel A. Antibodies against IL-1β, Caspase-1 (CASP1) and NLRP3 were used. An antibody against Tubulin was used as a loading control. (**C**) ELISA measurements of IL-1β in culture supernatants from LPS-primed THP-1 cells treated as indicated with 100 µM R837, 25 µM Yoda1 or 100 µM R837 plus 25 µM Yoda1 in presence of TRAM34, 10 µM MCC950 or extracellular KCl for 1 h. (**D**) Immunoblotting of culture supernatants (Sup) and lysates (Lys) from THP-1 cells in experiments as described for panel C. (**E**) Uptake of Sytox Green in THP-1 cells in experiments as described for panel C. (**F**) ELISA measurements of IL-1β in culture supernatants from LPS-primed BMDMs treated as indicated with 50 µM R837, 25 µM Yoda1 or 50 µM R837 plus 25 µM Yoda1 in presence of TRAM34, 10 µM MCC950 or extracellular KCl for 1 h. (**G**) Immunoblotting of culture supernatants (Sup) and lysates (Lys) from LPS-primed BMDMs treated as for panel F. Antibodies against IL-1β, Caspase-1 (CASP1) and NLRP3 were used. An antibody against Tubulin was used as a loading control. “ND” not detected. \**p < 0.05*, \*\**p < 0.01*, \*\*\**p < 0.001*. Data are representative of at least three independent experiments.

**Fig. S10.**
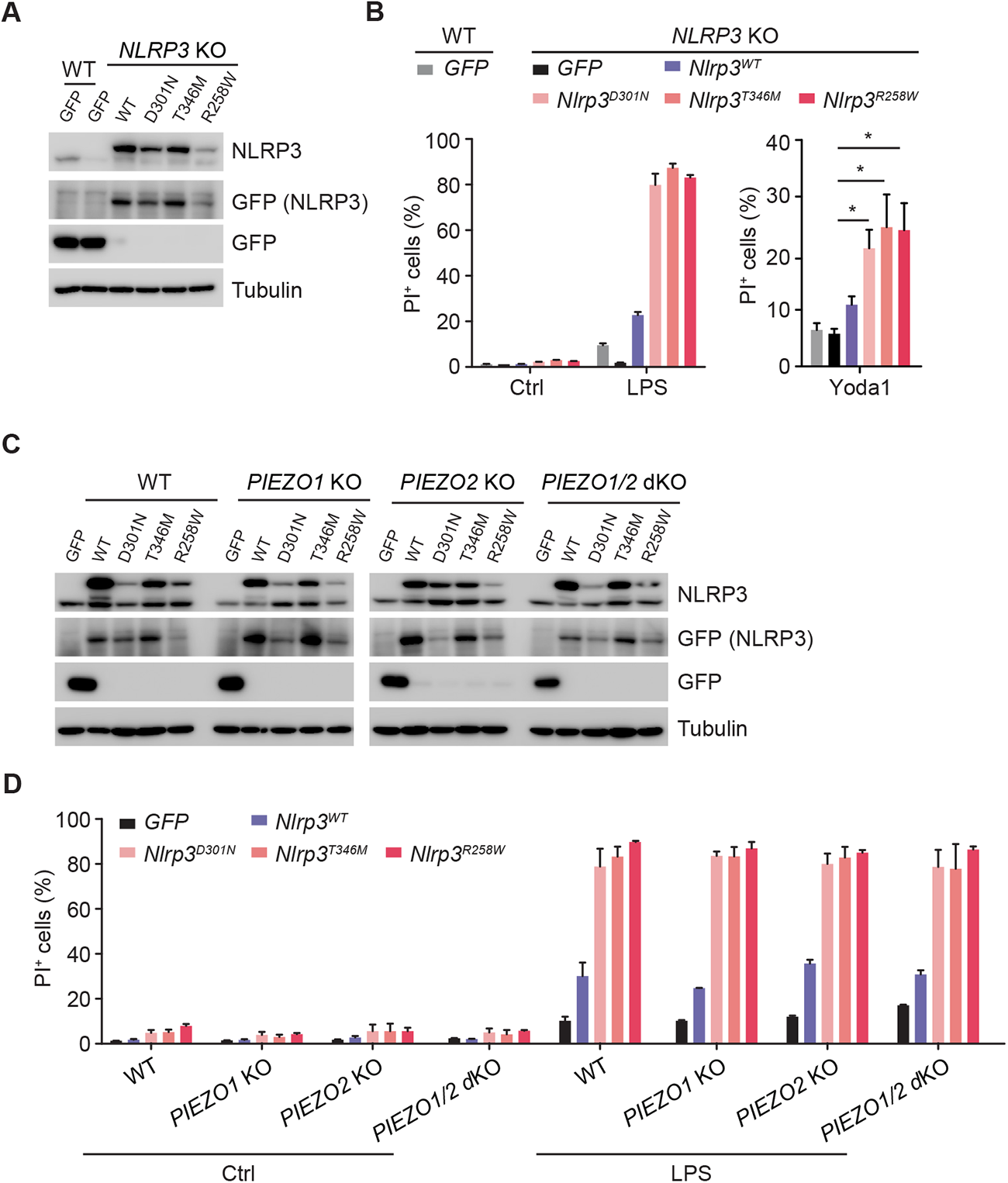
Yoda1 is sufficient to activate the NLRP3 inflammasome in cells expressing CAPS-causing NLRP3 mutants. (**A**) Immunoblotting of lysates from WT and *NLRP3* KO THP-1 cells expressing GFP, WT- or D301N-, T346M-, R258W mutated Nlrp3. Antibodies against NLRP3 and GFP were used. An antibody against Tubulin was used as a loading control. (**B**) Uptake of Propidium Iodide (PI) in WT and *NLRP3* KO THP-1 cells expressing GFP, WT or D301N-, T346M-, R258W-mutated mouse Nlrp3. Cells were treated with vehicle,1 µg/ml LPS or 25 μM Yoda1. Propidium Iodide uptake was analyzed by FACS after staining. (**C**) Immunoblotting of lysates from WT, *PIEZO1* KO, *PIEZO2* KO and *PIEZO1/2* dKO THP-1 cells expressing GFP, WT or D301N-, T346M-, R258W-mutated Nlrp3. Antibodies against NLRP3 and GFP were used. An antibody against Tubulin was used as a loading control. (**D**) Uptake of Propidium Iodide (PI) in WT, *PIEZO1* KO, *PIEZO2* KO and *PIEZO1/2* dKO THP-1 cells expressing GFP, WT or D301N-, T346M-, R258W-mutated mouse Nlrp3. Cells were treated with vehicle or 1 µg/ml LPS for 6 h. Propidium Iodide uptake was analyzed by FACS after staining. \**p < 0.05*. Data are representative of at least three independent experiments.

## Acknowledgments

We thank all the membranes in Ricci’s laboratory for scientific inputs. We also thank the facilities at IGBMC, including flow cytometry facility, cell culture facility, imaging facility and animal facility, for their technique help during the whole study.

## Author contributions

Conceptualization, ZZ and RR; Methodology, ZZ, LR and TY; Investigation, ZZ and LR; Writing – Review & Editing, RR, ZZ and LR; Funding Acquisition, RR, ZZ, NS, SE and IS; Resources, ZZ, LR, EE and SE; Supervision, ZZ, NS, SE and RR.

## Data and materials availability

All data is available in the main text or the supplementary materials. Requests of new materials created in this study should be sent to

